# Age-Dependent Spatiotemporal Remodeling of Brain Sphingolipids During LPS-Induced Neuroinflammation: MALDI-MSI Reveals Accelerated Sphingomyelin Depletion and Sulfatide Accumulation Linked to Mitochondrial Oxidative Stress

**DOI:** 10.64898/2026.06.26.734865

**Authors:** Kent Brown, Brayden Storey, Jacob Williams, Derrick Simet, Muhammad Bilal Umar, Emma Madsen, Zhiying Shan, Lanrong Bi

## Abstract

Aging is a major risk factor for exacerbated neuroinflammation and neurodegenerative diseases, yet the underlying lipid metabolic mechanisms remain incompletely understood. Here, we employed high-resolution matrix-assisted laser desorption/ionization mass spectrometry imaging (MALDI-MSI) combined with quantitative peak-area analysis and conceptual kinetic modeling to investigate age-dependent sphingolipid remodeling in the rat brain following intracerebro-ventricular (ICV) LPS challenge.

In old rats, MALDI-MSI revealed pronounced and progressive sphingolipid dysregulation compared with young animals. Quantitative analysis showed a dramatic ∼10-fold reduction in SM(d36:1) and ∼4-fold reduction in SM(d42:2), accompanied by significant accumulation of long-chain sulfatides (2.12-fold increase in C24:1-sulfatide and 1.45-fold increase in C24(OH)-sulfatide) at both 24 h and 72 h post-LPS. Spatial imaging demonstrated that these changes were markedly amplified in white matter regions and became more widespread and intense at 72 h.

A simplified Michaelis-Menten kinetic model successfully recapitulated the experimental data, identifying increased nSMase2 activity (higher Vmax) as the primary driver of accelerated sphingomyelin hydrolysis and subsequent ceramide rerouting into sulfatide synthesis. This metabolic shift generates excess ceramide that promotes Drp1-mediated mitochondrial fission, elevates mitochondrial ROS production, and disrupts bioenergetics, establishing a feed-forward loop linking sphingolipid remodeling to mitochondrial oxidative stress and white matter vulnerability in the aged brain.

These findings provide the first spatially and temporally resolved demonstration of age-dependent sphingolipid metabolic reprogramming during neuroinflammation. By integrating multimodal MALDI-MSI, quantitative lipidomics, and kinetic modeling, this study reveals a previously underappreciated nSMase2-ceramide-mitochondrial axis in neuroinflammaging.

**Graphic Abstract:** 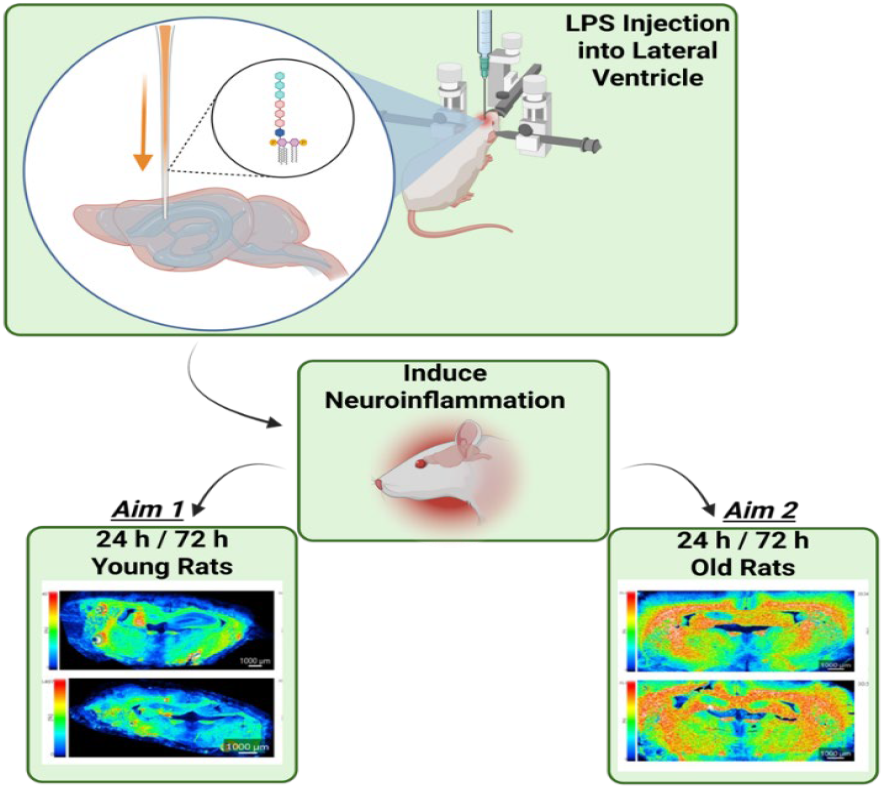

## 1. Introduction

Aging is accompanied by a chronic, low-grade, sterile inflammatory state known as “inflammaging,” a concept first proposed by Franceschi et al. in 2000 [1]. This condition arises from the lifelong accumulation of molecular and cellular damage, which leads to persistent activation of the innate immune system even in the absence of infection [2,3]. In the central nervous system (CNS), inflammaging manifests as neuroinflammaging, characterized by sustained microglial and astrocytic activation, elevated production of pro-inflammatory cytokines (such as IL-6, TNF-α, and IL-1β), increased oxidative stress, and impaired resolution of inflammatory responses [4].

A key feature of neuroinflammaging is the primed phenotype of aged microglia. These cells exhibit heightened reactivity to secondary inflammatory challenges, such as lipopolysaccharide (LPS), infection, or damage-associated molecular patterns (DAMPs). This priming results in exaggerated and prolonged neuroinflammatory responses in aged individuals compared with younger ones [5]. In addition, the accumulation of senescent cells in the aged brain contributes to the inflammatory environment through the senescence-associated secretory phenotype (SASP), which involves the release of pro-inflammatory cytokines, chemokines, and matrix-remodeling factors [6,7]. Together, these changes create a basal inflammatory tone that disrupts brain homeostasis and increases susceptibility to neurodegenerative processes [8].

Neuroinflammaging has emerged as a major contributor to the pathogenesis of age-related neurodegenerative diseases, including Alzheimer’s disease (AD) and Parkinson’s disease (PD) [4]. Chronic glial activation and dysregulated cytokine signaling promote neuronal dysfunction, synaptic loss, and protein aggregation [9]. The aged brain’s heightened vulnerability to inflammatory stimuli, such as systemic or intracerebro-ventricular (ICV) LPS administration, therefore provides a valuable experimental model to investigate how aging modifies lipid metabolism, membrane integrity, and inflammatory signaling [10,11].

Sphingolipids are highly enriched in the CNS and serve both structural and signaling functions. In myelin, galactosylceramide (GalCer) and its sulfated derivative, sulfatide, account for approximately 25% of total myelin lipids [12,13]. These lipids are essential for myelin formation, compaction, and long-term stability, contributing to the unique biophysical properties required for rapid saltatory conduction [14]. Genetic deficiencies in enzymes involved in GalCer or sulfatide synthesis result in severe myelin abnormalities and neurological dysfunction, underscoring their structural importance [15,16].

Beyond their role in myelin, sphingolipids are critical organizers of lipid rafts and cholesterol- and sphingolipid-enriched membrane microdomains that compartmentalize signaling molecules [17]. Sphingomyelin, ceramide, and glycosphingolipids within these rafts facilitate the spatial organization of receptors and signaling proteins in neurons, oligodendrocytes, and astrocytes [18]. Ceramide, in particular, can self-associate to form large membrane platforms that promote receptor clustering and the activation of pro-apoptotic and pro-inflammatory pathways [19]. In contrast, sphingosine-1-phosphate (S1P), derived from ceramide, regulates cell survival, migration, and inflammation through G-protein-coupled receptors [20]. Thus, sphingolipids serve dual roles as structural components of myelin and as dynamic regulators of membrane signaling and cellular responses to stress.

Neutral sphingomyelinase 2 (nSMase2), encoded by SMPD3, is the predominant neutral sphingomyelinase in the brain and functions as a critical link between inflammatory signaling and ceramide production. Unlike acid sphingomyelinase, which resides in lysosomes, nSMase2 is localized to the plasma membrane and Golgi apparatus. It hydrolyzes sphingomyelin to generate ceramide and phosphocholine in response to stress and inflammatory stimuli [21]. Activation of nSMase2 occurs rapidly downstream of Toll-like receptor 4 (TLR4) following LPS exposure, as well as through TNF-α, IL-1β, and oxidative stress pathways [22]. Upon stimulation, nSMase2 translocates to the plasma membrane, where it locally produces ceramide that reorganizes membrane microdomains into platforms, facilitating receptor clustering and activation of downstream inflammatory cascades such as NF-κB and MAPK signaling [23].

The resulting increase in ceramide promotes microglial activation, cytokine production, exosome biogenesis, and, in certain contexts, neuronal apoptosis and mitochondrial dysfunction [24,25,26]. Importantly, nSMase2 activity and expression are elevated in the aged brain, contributing to the heightened inflammatory tone of neuroinflammaging [27]. Pharmacological or genetic inhibition of nSMase2 has been shown to attenuate LPS-induced microglial activation and neuroinflammation, underscoring its role in amplifying inflammatory responses [28].

Despite substantial evidence that neuroinflammaging increases brain vulnerability to inflammatory insults, significant gaps remain in our understanding of how aging alters sphingolipid metabolism during acute inflammation. Most previous studies have relied on bulk lipidomic analyses, which lack spatial resolution, or have examined sphingolipid changes in the context of normal aging or chronic disease, with limited direct comparison between young and aged brains after a controlled inflammatory challenge such as LPS [29,30]. Furthermore, few studies have utilized high-resolution matrix-assisted laser desorption/ionization mass spectrometry imaging (MALDI-MSI) to simultaneously capture both the spatial distribution and relative quantitative abundance of sphingolipids within the same tissue sections [31,32].

To address these limitations, our present study employed high-resolution MALDI-MSI using the iMScope QT instrument to investigate sphingolipid remodeling in young and aged rat brains at 24 h and 72 h after intracerebroventricular LPS administration. This approach enabled both spatial mapping of sulfatide and sphingomyelin species across brain regions and relative quantification through peak area analysis from the same dataset. The specific objectives were to characterize spatial and quantitative changes in key sphingolipids associated with aging and inflammation, to integrate spatial distribution patterns with quantitative data for a comprehensive understanding of regional lipid remodeling, and to explore potential mechanistic links between observed lipid changes and enzymes such as nSMase2 that mediate sphingomyelin hydrolysis and ceramide rerouting. By applying high-resolution MALDI-MSI in a young-versus-aged LPS model, this study aims to provide new insights into the sphingolipid alterations that accompany neuroinflammaging and to establish a foundation for future investigations into their functional consequences.

## 2. Experimental Section

### 2.1 Animals and LPS-induced Neuroinflammation Model

All animal procedures were approved by the Michigan Technological University Institutional Animal Care and Use Committee (IACUC) and performed in accordance with the National Institutes of Health Guide for the Care and Use of Laboratory Animals. Adult Sprague Dawley (SD) rats were purchased from Charles River Laboratories (Wilmington, MA) and housed in the Michigan Tech Animal Care Facility (ACF). Upon arrival, all rats underwent a 1–2-week acclimatization period to allow adjustment to the housing environment prior to the initiation of experimental procedures. Neuroinflammation was induced by stereotaxic administration of lipopolysaccharide (LPS) into the right lateral ventricle. At 24 h and 72 h post-LPS administration, rats were euthanized and brain tissues were rapidly excised for matrix-assisted laser desorption/ionization mass spectrometry imaging (MALDI-MSI) analysis.

### 2.2. Tissue Processing for MALDI-MSI

Following excision, all brain tissues were inactivated by treatment with 70% ethanol prior to cryo-sectioning and mounting onto indium tin oxide (ITO)-coated slides. This chemical inactivation method was selected because it is minimally invasive while remaining compatible with downstream lipid and metabolite imaging.

### 2.3. MALDI-MSI Parameters

MALDI mass spectrometry imaging was performed using the 5 µm method with the parameters listed in Table 1.

## 3. Results

### 3.1 Quantitative changes in sulfatide levels

Quantitative analysis of sulfatide species revealed prominent age-associated differences following LPS-induced neuroinflammation. Two-way ANOVA demonstrated a significant main effect of age on the relative abundance of C24:1-sulfatide (F(1,2) = 67.90, p = 0.014), with old rats exhibiting substantially higher levels than young rats at both 24 h and 72 h post-LPS administration. Post-hoc comparisons indicated that C24:1-sulfatide levels were elevated by approximately 2.12-fold in old rats compared with young animals. A comparable but more modest age-related increase was observed for C24(OH)-sulfatide, which showed an approximate 1.45-fold elevation in the old group. Although the Age × Time interaction did not reach statistical significance, the magnitude of the age-associated increase in both C24 sulfatide species tended to be greater at 72 h than at 24 h, suggesting progressive accumulation in the aged brain following inflammatory challenge.

**Table 1.** MALDI-MSI Parameters.

| Parameter | Value | Unit / Notes |
| --- | --- | --- |
| Method | 5 μm | — |
| Pitch | 25 | μm (step size) |
| Laser shots | 80 | per pixel |
| Laser repetition rate | 2000 | Hz |
| Laser diameter | 2 | (as provided) |
| Laser intensity | 65 | (arbitrary units) |
| Polarity | Positive or Negative | Both modes acquired |
| Event type | MS | — |
| Measurement mode | Normal | — |
| Mass range | 500–1200 | m/z |
| Detector voltage | 2.38 | kV |
| Desolvation line (DL) temperature | 250 | °C |
| Heat block temperature | 450 | °C |

These quantitative findings indicate that aging markedly enhances the accumulation of long-chain sulfatides in response to an acute neuroinflammatory stimulus. The approximately 2-fold increase in C24:1-sulfatide and the more modest but consistent elevation in C24(OH)-sulfatide suggest that aging promotes both increased synthesis and/or reduced clearance of these myelin-enriched lipids. The trend toward greater accumulation at 72 h compared with 24 h further implies that the effects of aging on sulfatide metabolism are time-dependent, with prolonged inflammatory exposure leading to more pronounced lipid dysregulation in old animals. These changes may reflect heightened nSMase2 activity in the aged brain, which accelerates sphingomyelin hydrolysis and increases ceramide availability for downstream conversion into sulfatides. The observed age-dependent elevation in sulfatides is therefore consistent with a model in which aging amplifies inflammation-driven sphingolipid remodeling, potentially contributing to altered myelin composition and membrane properties in the aged brain. These quantitative differences, derived from peak area measurements obtained through high-resolution MALDI-MSI, are summarized in Table 2.

### 3.2 Spatial distribution of sulfatides revealed by MALDI-MSI

MALDI-MSI analysis revealed clear age- and time-dependent differences in the spatial distribution of C24 sulfatides following LPS challenge. In young rats, both C24:1-sulfatide (m/z 888.624) and C24(OH)-sulfatide (m/z 906.635) exhibited moderate signal intensity with relatively localized and structured distribution at both 24 h and 72 h post-LPS (Figure 1A, Panels A and C; Figure 1B, Panels E and G). High-intensity regions largely respected anatomical boundaries.

**Figure 1A.**
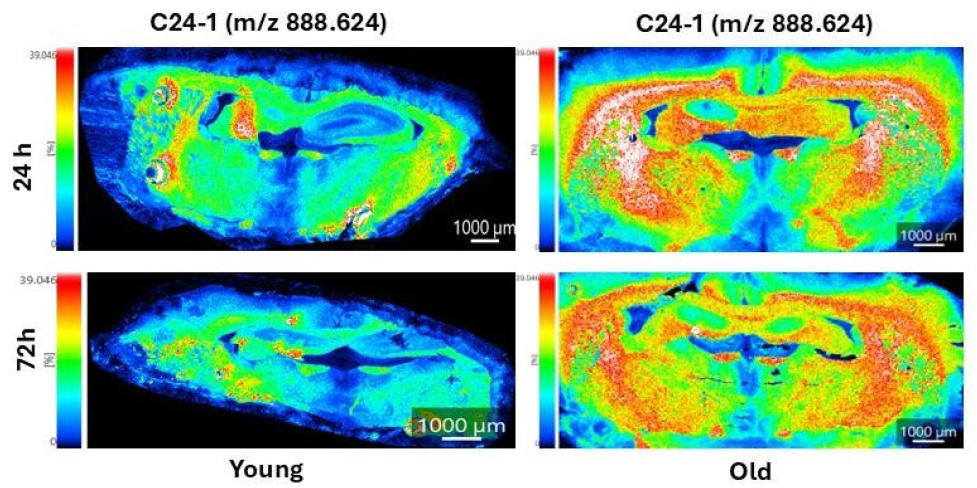
Spatial distribution of C24:1-sulfatide in young and old rat brains at 24 h and 72 h after LPS challenge. Representative MALDI-MSI ion images showing the spatial distribution and relative abundance of C24:1-sulfatide (m/z 888.624) in young and old rat brain sections at 24 h and 72 h following intracerebroventricular LPS injection. Images are arranged for direct age and time comparison. In young rats, C24:1-sulfatide signal remained relatively structured and regionally confined, with high-intensity areas largely respecting anatomical boundaries at both time points. In contrast, old rats exhibited markedly higher signal intensity and a more diffuse, less spatially restricted distribution. This morphological shift from localized to widespread and scattered distribution was already detectable at 24 h but became dramatically more pronounced at 72 h, with strong accumulation extending across large areas, particularly in white matter regions. All images are displayed on the same normalized intensity scale. Scale bar = 1000 μm.

**Figure 1B.**
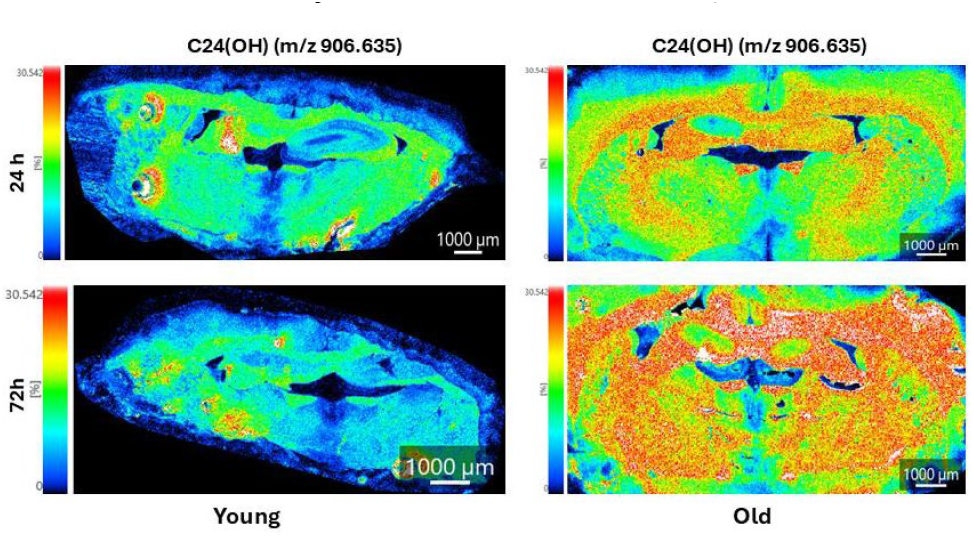
Spatial distribution of C24(OH)-sulfatide in young and old rat brains at 24 h and 72 h after LPS challenge. Representative MALDI-MSI ion images showing the spatial distribution and relative abundance of C24(OH)-sulfatide (m/z 906.635) in young and old rat brain sections at 24 h and 72 h following intracerebroventricular LPS injection. Images are arranged for direct age and time comparison. In young rats, C24(OH)-sulfatide signal remained relatively structured and regionally confined at both 24 h and 72 h, with moderate increases in intensity over time. In contrast, old rats exhibited markedly higher signal intensity and a highly diffuse, less spatially restricted distribution. This morphological shift was already evident at 24 h but became dramatically more pronounced at 72 h, where the signal appeared highly scattered and “cloud-like,” extending across large areas of the brain section with minimal respect for anatomical boundaries. The loss of spatial organization was more severe for C24(OH)-sulfatide than for C24:1-sulfatide, with particularly strong accumulation in myelinated white matter regions. All images are displayed on the same normalized intensity scale. Scale bar = 1000 μm.

In contrast, old rats displayed markedly higher signal intensity and more widespread distribution of both sulfatide species. These age-related differences were already detectable at 24 h but became substantially more pronounced at 72 h, where intense signals extended across large areas of the brain sections (Figure 1A, Panel D; Figure 1B, Panel H). The most striking spatial changes were observed for C24(OH)-sulfatide in old rats at 72 h.

Notably, MALDI-MSI also revealed distinct morphological differences in sulfatide distribution between age groups. In young rats, the signal remained relatively structured and regionally confined at both time points. In old rats, however, the distribution shifted from moderately localized at 24 h to highly diffuse and less anatomically restricted by 72 h, appearing scattered and “cloud-like” across broad regions. This morphological transition from localized to diffuse distribution was particularly evident for C24(OH)-sulfatide and indicates that aging impairs the brain’s ability to maintain spatial containment of sulfatide accumulation during prolonged neuroinflammation.

**Table 2.** Quantitative analysis of sulfatide levels in young and old rat brains following LPS challenge.

| Sulfatide Species | m/z | Young<br>(Mean ± SD) | Old<br>(Mean ± SD) | Fold Change<br>(Old/Young) | Age Effect<br>(p-value) | Notes |
| --- | --- | --- | --- | --- | --- | --- |
| C24:1-Sulfatide | 888.624 | 90,139 ± 20,578 | 191,139 ± 5,845 | 2.12 | 0.014 | Significant main effect of Age |
| C24(OH)-Sulfatide | 906.635 | ~71,000 | ~103,000 | ~1.45 | n.s. | Moderate increase in Old rats |
| C22(OH)-Sulfatide | 878.603 | ~30,000 | ~53,000 | ~1.77 | n.s. | Trend toward increase in Old |
| C18:1-Sulfatide | 806.546 | 33,734 ± 8,681 | 32,016 ± 1,639 | 0.95 | n.s. | No significant change |

Regional quantification further demonstrated that the age-related increase in sulfatides was not uniform. The strongest elevation of C24(OH)-sulfatide occurred in white matter regions such as the corpus callosum (Supplementary Table S1). This regional preference is consistent with the high baseline sulfatide content of myelin and suggests that myelinated white matter is particularly susceptible to inflammation-driven sphingolipid remodeling in the aged brain.

These spatial alterations were consistent with the quantitative increases detected by peak area analysis (Table 2), confirming that the age-dependent elevation in sulfatides represents both a quantitative and spatially heterogeneous phenomenon. These findings indicate that aging modifies not only the abundance but also the regional distribution and morphological organization of long-chain sulfatides during neuroinflammation, with pronounced effects in white matter.

### 3.3 Time-dependent changes in sulfatide distribution in young rats

MALDI-MSI analysis revealed progressive but moderate and spatially controlled changes in sulfatide distribution in young rat brains following LPS challenge. As shown in Figure 2A, signal intensity for C18-1-sulfatide (m/z 806.546), C18(OH)-sulfatide (m/z 822.541), C22-sulfatide (m/z 862.608), and C22(OH)-sulfatide (m/z 878.603) was generally moderate at 24 h post-LPS. By 72 h, several species—particularly the hydroxylated forms—exhibited noticeable increases in signal intensity. These increases were accompanied by modest expansion in regional coverage; however, the overall distribution remained relatively structured and regionally defined, with high-intensity areas largely respecting anatomical boundaries. This pattern suggests that young animals respond to LPS challenge with a progressive yet spatially regulated sulfatide remodeling process.

**Figure 2A.**
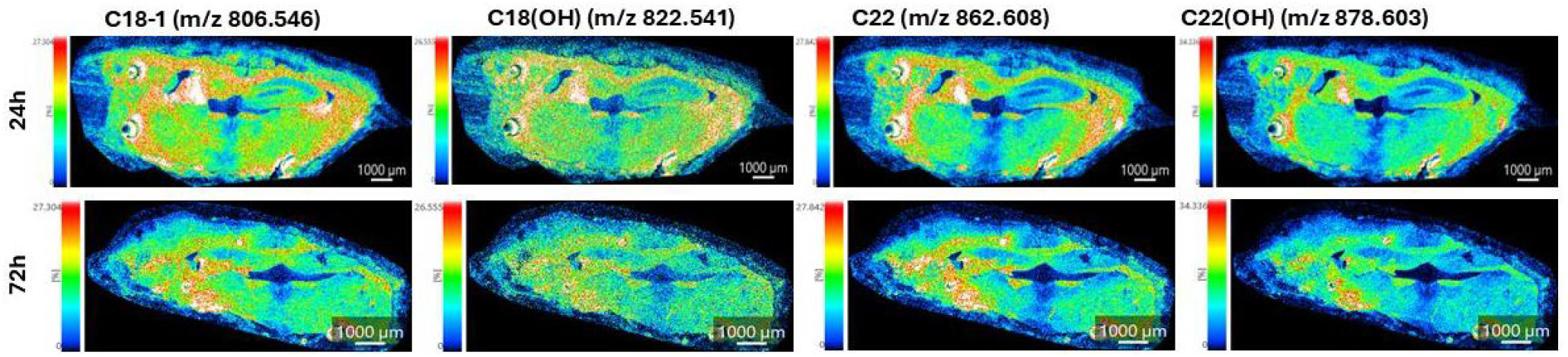
Time-dependent changes in C18 and C22 sulfatide distribution in young rat brain after LPS challenge. Representative MALDI-MSI ion images showing the spatial distribution and relative abundance of C18-1-sulfatide (m/z 806.546), C18(OH)-sulfatide (m/z 822.541), C22-sulfatide (m/z 862.608), and C22(OH)-sulfatide (m/z 878.603) in young rat brain sections at 24 h and 72 h following intracerebroventricular LPS injection. In young rats, LPS induced progressive but moderate and relatively controlled increases in sulfatide signal intensity from 24 h to 72 h. At 24 h, the signals were moderate in intensity and spatially structured, with high-intensity regions largely respecting anatomical boundaries. By 72 h, several species—particularly the hydroxylated forms (C18(OH)-sulfatide and C22(OH)-sulfatide)—showed noticeable increases in signal intensity; however, the distribution remained organized and regionally defined rather than becoming diffusely widespread. This morphological pattern indicates that young animals exhibit a more contained and spatially regulated sulfatide remodeling response during neuroinflammation. All images within each sulfatide species are displayed on the same normalized intensity scale. Scale bar = 1000 μm.

**Figure 2B.**
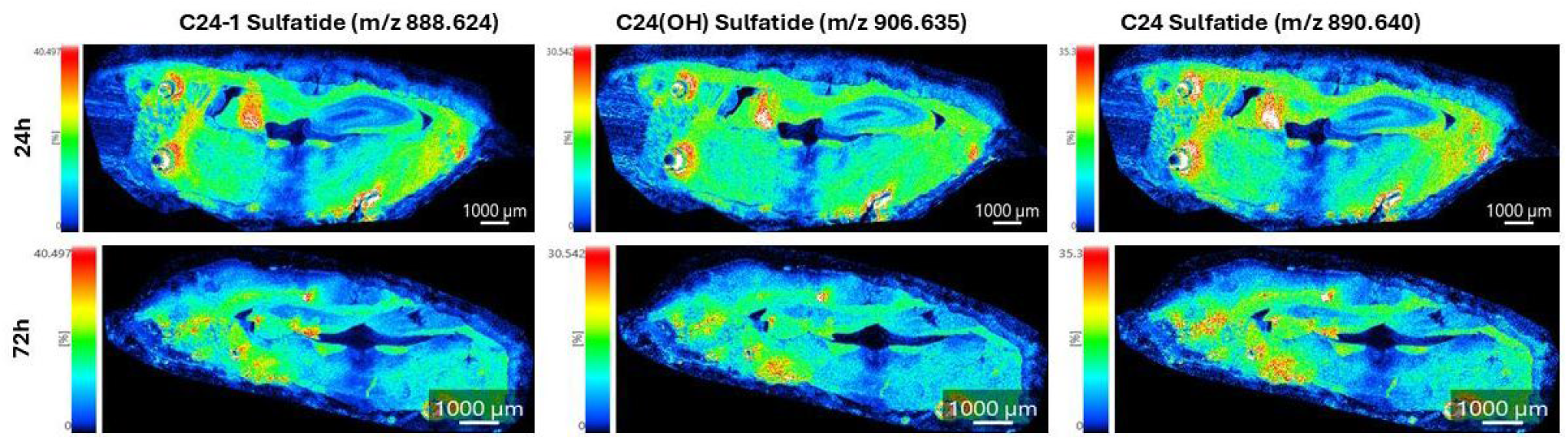
Time-dependent changes in C24 sulfatide distribution in young rat brain after LPS challenge. Representative MALDI-MSI ion images showing the spatial distribution and relative abundance of C24 sulfatide species, including C24:1-sulfatide (m/z 888.624), C24(OH)-sulfatide (m/z 906.635), and C24-sulfatide (m/z 890.640), in young rat brain sections at 24 h and 72 h following intracerebroventricular LPS injection. In young rats, LPS induced progressive but controlled increases in sulfatide signal intensity from 24 h to 72 h. At 24 h, C24-1-sulfatide, C24(OH)-sulfatide, and C24-sulfatide exhibited moderate signal intensity with relatively structured and regionally defined distribution. By 72 h, all three species showed noticeable increases in signal intensity, particularly C24:1-sulfatide and C24(OH)-sulfatide; however, the signals remained spatially organized and largely confined to specific anatomical regions rather than becoming diffusely widespread. These morphological patterns indicate that young animals mount a more contained and spatially regulated sulfatide remodeling response during neuroinflammation, in contrast to the highly diffuse and less anatomically restricted accumulation observed in aged rats. All images within each sulfatide species are displayed on the same normalized intensity scale to facilitate direct temporal comparison. Scale bar = 1000 μm.

In the C24 sulfatide group (Figure 2B), more pronounced time-dependent changes were observed compared with C18 and C22 species. C24:1-sulfatide (m/z 888.624) and C24(OH)-sulfatide (m/z 906.635) showed clear increases in both signal intensity and regional coverage from 24 h to 72 h, whereas C24-sulfatide (m/z 890.640) displayed a more modest increase. Notably, the hydroxylated C24(OH)-sulfatide exhibited the most substantial rise in intensity among the C24 species, suggesting that hydroxylation may enhance the responsiveness of long-chain sulfatides to inflammatory stimuli. Despite these elevations, the sulfatide signals in young rats remained spatially organized and largely confined to specific anatomical regions rather than becoming diffusely widespread.

These observations indicate species- and chain-length-dependent differences in sulfatide remodeling in young rats. Longer-chain C24 sulfatides underwent greater changes in both abundance and spatial coverage compared with shorter-chain C18 and C22 species, consistent with preferential metabolic routing of longer-chain ceramides into the sulfatide biosynthetic pathway during neuroinflammation. The retention of structured and regionally defined distribution at 72 h, even with increased signal intensity, suggests that young animals maintain effective spatial and temporal regulation of sulfatide changes. This controlled remodeling response stands in clear contrast to the highly diffuse and less anatomically restricted accumulation observed in aged rats, highlighting fundamental age-dependent differences in the brain’s ability to regulate sphingolipid metabolism during inflammatory challenge.

### 3.4 Time-dependent changes in sulfatide distribution in old rats

MALDI-MSI demonstrated robust and progressive increases in sulfatide distribution in old rats from 24 h to 72 h post-LPS (Figure 3A and Figure 3B). Both C18/C22 and C24 sulfatide species exhibited marked elevations in signal intensity and spatial spread, with the most striking increases observed in C24:1-sulfatide and C24(OH)-sulfatide at 72 h. These changes were substantially greater in both magnitude and spatial extent than those observed in young rats over the same period (Figures 2A and 2B). While young rats showed moderate increases in signal intensity with relatively structured and regionally confined distribution, old rats displayed highly diffuse and less anatomically restricted patterns, particularly at 72 h, where the signals appeared scattered and “cloud-like” across large areas of the brain sections.

**Figure 3A.**
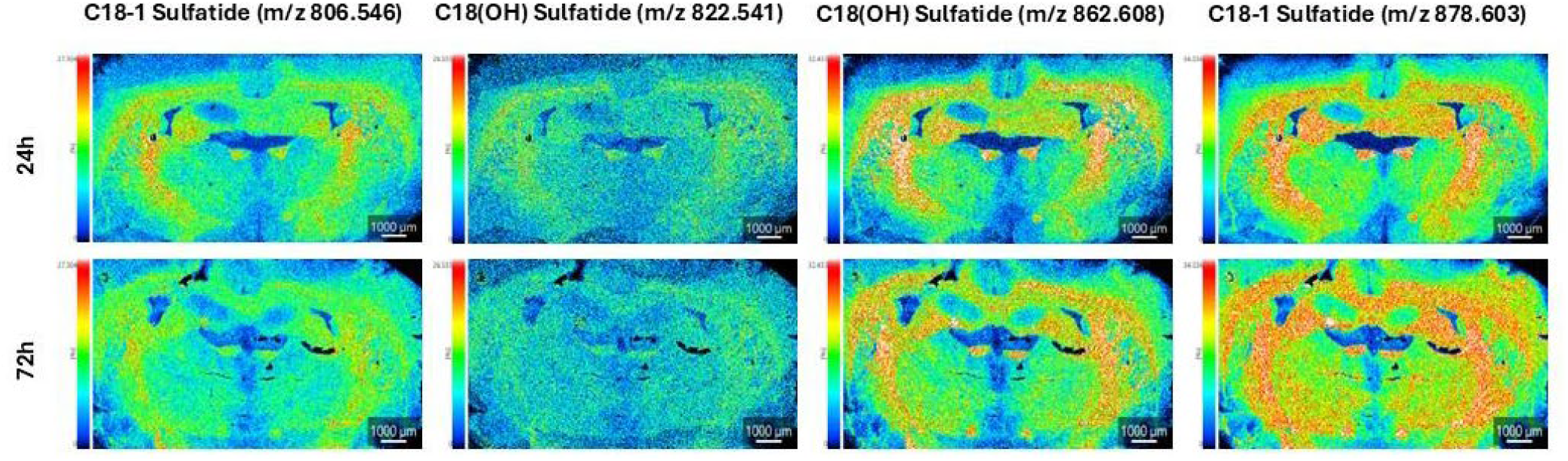
Time-dependent changes in C18 and C22 sulfatide distribution in old rat brain after LPS challenge. Representative MALDI-MSI ion images showing the spatial distribution and relative abundance of C18-1-sulfatide (m/z 806.546), C18(OH)-sulfatide (m/z 822.541), C22-sulfatide (m/z 862.608), and C22(OH)-sulfatide (m/z 878.603) in old rat brain sections at 24 h and 72 h following intracerebroventricular LPS injection. In old rats, LPS induced robust and progressive increases in sulfatide signal intensity and spatial spread from 24 h to 72 h. The increases were particularly pronounced in white matter regions, with signals becoming markedly more intense and widespread at the later time point. In contrast to the relatively structured and regionally confined distribution observed in young rats, old rats exhibited a highly diffuse and less anatomically restricted pattern, especially at 72 h. All images within each sulfatide species are displayed on the same normalized intensity scale to enable direct temporal comparison. Scale bar = 1000 μm.

**Figure 3B.**
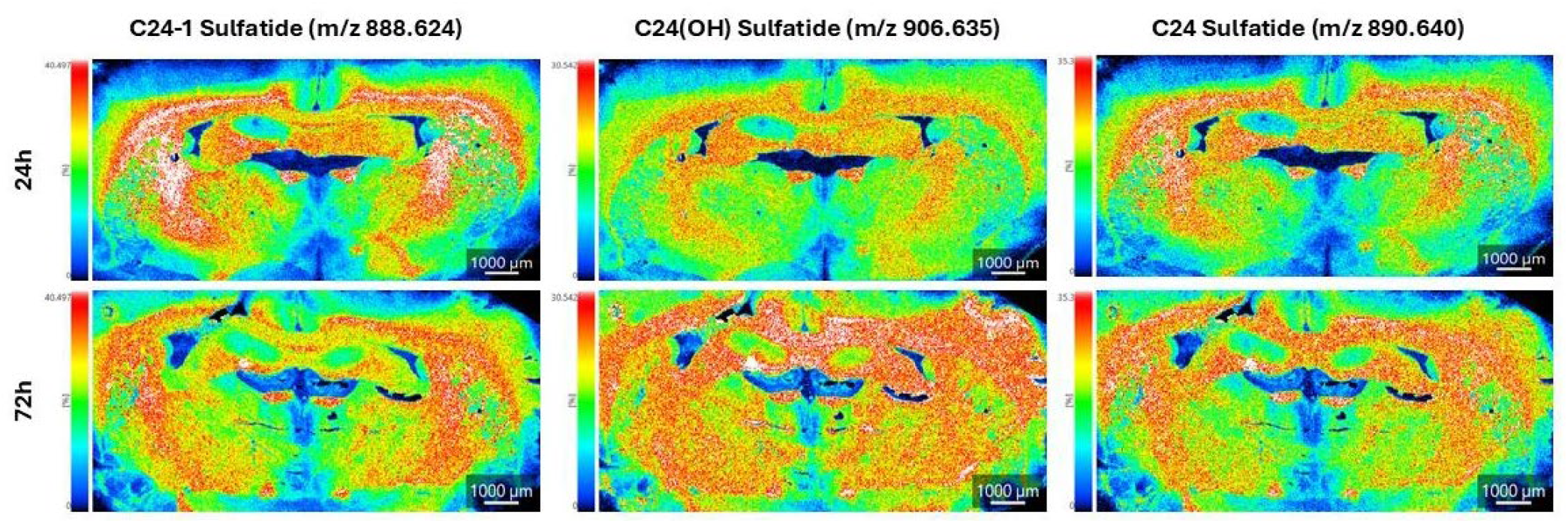
Time-dependent changes in C24 sulfatide distribution in old rat brain after LPS challenge. Representative MALDI-MSI ion images showing the spatial distribution and relative abundance of C24 sulfatide species, including C24:1-sulfatide (m/z 888.624), C24(OH)-sulfatide (m/z 906.635), and C24-sulfatide (m/z 890.640), in old rat brain sections at 24 h and 72 h following intracerebroventricular LPS injection. In old rats, LPS induced robust and progressive increases in both signal intensity and spatial spread from 24 h to 72 h. At 24 h, all three C24 sulfatide species already exhibited high signal intensity with widespread distribution. By 72 h, the signals became markedly more intense and highly diffuse, appearing scattered and less anatomically restricted across large areas of the brain sections. This morphological transition from widespread to extremely diffuse distribution was especially pronounced in white matter regions and was more severe than the relatively structured and regionally confined changes observed in young rats. All images within each sulfatide species are displayed on the same normalized intensity scale to enable direct temporal comparison. Scale bar = 1000 μm.

These robust time-dependent increases indicate that aging substantially amplifies both the rate and extent of sulfatide remodeling following an acute inflammatory challenge. The pronounced elevation in C24 species, especially C24(OH)-sulfatide, suggests that longer-chain and hydroxylated sulfatides are preferentially affected in the aged brain. This may reflect differences in substrate preference of sulfatide-synthesizing enzymes (UGT8 and CST) or greater metabolic stability of these species under sustained inflammatory conditions. In contrast to the controlled and spatially organized remodeling seen in young rats, the distribution in old rats became progressively more diffuse and widespread from 24 h to 72 h, indicating a loss of spatial containment.

The progressive nature of these changes is consistent with sustained nSMase2 activity in old rats, leading to continued sphingomyelin hydrolysis and cumulative ceramide production that is channeled into sulfatide synthesis over time. The rapid and extensive sulfatide accumulation observed in old rats at 72 h, particularly in white matter regions, suggests that aging impairs the brain’s ability to resolve or spatially restrict inflammation-associated lipid changes. This results in a more widespread and persistent alteration of sphingolipid composition, potentially contributing to greater disruption of myelin integrity and heightened local mitochondrial oxidative stress in aged white matter compared with the more contained response in young animals.

### 3.5 Quantitative and spatial alterations in sphingomyelins

In contrast to the progressive accumulation of sulfatides, sphingomyelin levels were substantially lower in old rats compared with young animals following LPS challenge. Quantitative peak-area analysis revealed a striking, species-specific depletion of sphingomyelin. SM(d36:1) exhibited an approximately 10-fold decrease in relative abundance in old rats, whereas SM(d42:2) showed a more moderate ∼4-fold reduction. This preferential loss of shorter-chain sphingomyelin suggests that SM(d36:1) may be more susceptible to enzymatic hydrolysis or resides in membrane environments that are more accessible to sphingomyelinases under inflammatory conditions in the aged brain.

These quantitative reductions were accompanied by marked spatial alterations revealed by MALDI-MSI. In young rats, sphingomyelin signals remained relatively structured and well-defined across brain sections at both 24 h and 72 h post-LPS. In old rats, however, sphingomyelin distribution was characterized by markedly reduced overall signal intensity and a more fragmented, diffuse pattern, particularly evident at 72 h. This morphological transition from organized to fragmented distribution indicates that sphingomyelin depletion in aged animals is accompanied by disruption of normal membrane microdomain organization and lipid raft integrity.

These quantitative and spatial findings support a model of enhanced nSMase2-mediated sphingomyelin hydrolysis in the aged brain during neuroinflammation. The substantial and preferential depletion of SM(d36:1), concurrent with the marked accumulation of long-chain sulfatides observed in Figures 1–4 and Table 2, points to a coordinated rerouting of sphingolipid metabolism. Increased nSMase2 activity accelerates sphingomyelin breakdown, generates a transient but significant ceramide pool, and redirects metabolic flux toward sulfatide synthesis. The resulting elevation in ceramide likely contributes to mitochondrial oxidative stress through Drp1-mediated fission and elevated mtROS production. The more pronounced sphingomyelin depletion and spatial fragmentation at 72 h further suggest that these effects are cumulative and become increasingly evident with prolonged inflammatory exposure in the aged brain, thereby exacerbating white matter vulnerability.

### 3.6 Differential remodeling between lipid classes and time points

Examination of lipid changes across classes revealed distinct and opposing patterns that were strongly influenced by age. While long-chain sulfatides (particularly C24:1-sulfatide and C24(OH)-sulfatide) showed significant increases in old rats following LPS administration (Figures 1B, 3B, 4), sphingomyelin species exhibited the opposite response, with substantial reductions, most notably for SM(d36:1) (Table 2). These divergent directional changes between sulfatides and sphingomyelins were consistently observed at both 24 h and 72 h post-LPS. Although the Age × Time interaction did not reach statistical significance, the magnitude of age-related alterations in both lipid classes was clearly greater at 72 h than at 24 h, indicating that the effects of aging on sphingolipid remodeling are progressive and become more pronounced with prolonged inflammatory exposure.

These findings demonstrate that aging alters both the direction and the temporal dynamics of sphingolipid remodeling following an inflammatory challenge. The inverse relationship between elevated sulfatide levels and reduced sphingomyelin levels in old rats is consistent with enhanced metabolic flux through the sphingomyelin-to-ceramide-to-sulfatide pathway. This pattern strongly suggests that aging promotes increased hydrolysis of sphingomyelin, most likely via elevated nSMase2 activity, resulting in greater ceramide availability that is subsequently utilized for sulfatide synthesis. The observation that these opposing changes become substantially more pronounced at 72 h compared with 24 h further implies that the effects of aging on this metabolic rerouting are cumulative over time.

Spatial imaging data reinforce and extend this interpretation. The most marked divergence between sulfatide accumulation and sphingomyelin depletion in old rats occurred at the later (72 h) time point (Figure 4), when inflammation-driven lipid remodeling had more time to progress. While young rats maintained relatively structured and controlled sulfatide distribution with only modest sphingomyelin changes over time (Figures 2A and 2B), old rats exhibited highly diffuse sulfatide accumulation alongside fragmented and reduced sphingomyelin signals (Figures 3A, 3B, and S1). These spatially resolved patterns indicate that aging not only shifts the balance between sphingomyelin and sulfatide but also impairs the brain’s ability to spatially contain and resolve these lipid alterations during sustained neuroinflammation. Collectively, the integration of quantitative (Table 2) and spatial data (Figures 1–4) demonstrates that aging accelerates and amplifies sphingolipid metabolic rerouting, resulting in coordinated sphingomyelin depletion and sulfatide accumulation that is both temporally progressive and spatially widespread, particularly in white matter regions.

**Figure 4.**
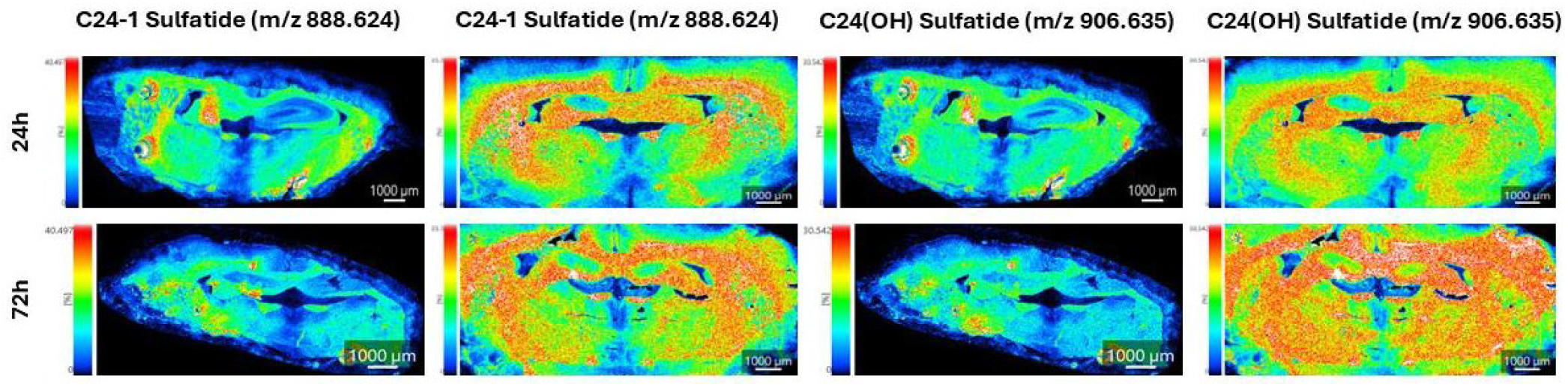
Age-dependent differences in C24 sulfatide distribution in young and old rat brains at 24 h and 72 h after LPS challenge. Representative MALDI-MSI ion images showing the spatial distribution and relative abundance of long-chain C24 sulfatide species (C24:1-sulfatide, m/z 888.624; and C24(OH)-sulfatide, m/z 906.635) in young and old rat brain sections at 24 h and 72 h following intracerebroventricular LPS injection. Images are arranged for direct visual comparison between age groups and time points. In young rats, both C24:1-sulfatide and C24(OH)-sulfatide signals remained relatively structured and regionally confined at both 24 h and 72 h, with moderate increases in intensity over time. In contrast, old rats already displayed high-intensity and widespread distribution at 24 h, which progressed to a highly diffuse and less anatomically restricted pattern by 72 h. This morphological transition from organized to scattered and “cloud-like” distribution was particularly pronounced for C24(OH)-sulfatide and was most evident in white matter-rich regions. All images are displayed on the same normalized intensity scale within each sulfatide species to allow accurate quantitative and spatial comparison. Scale bar = 1000 μm.

**Figure 5.**
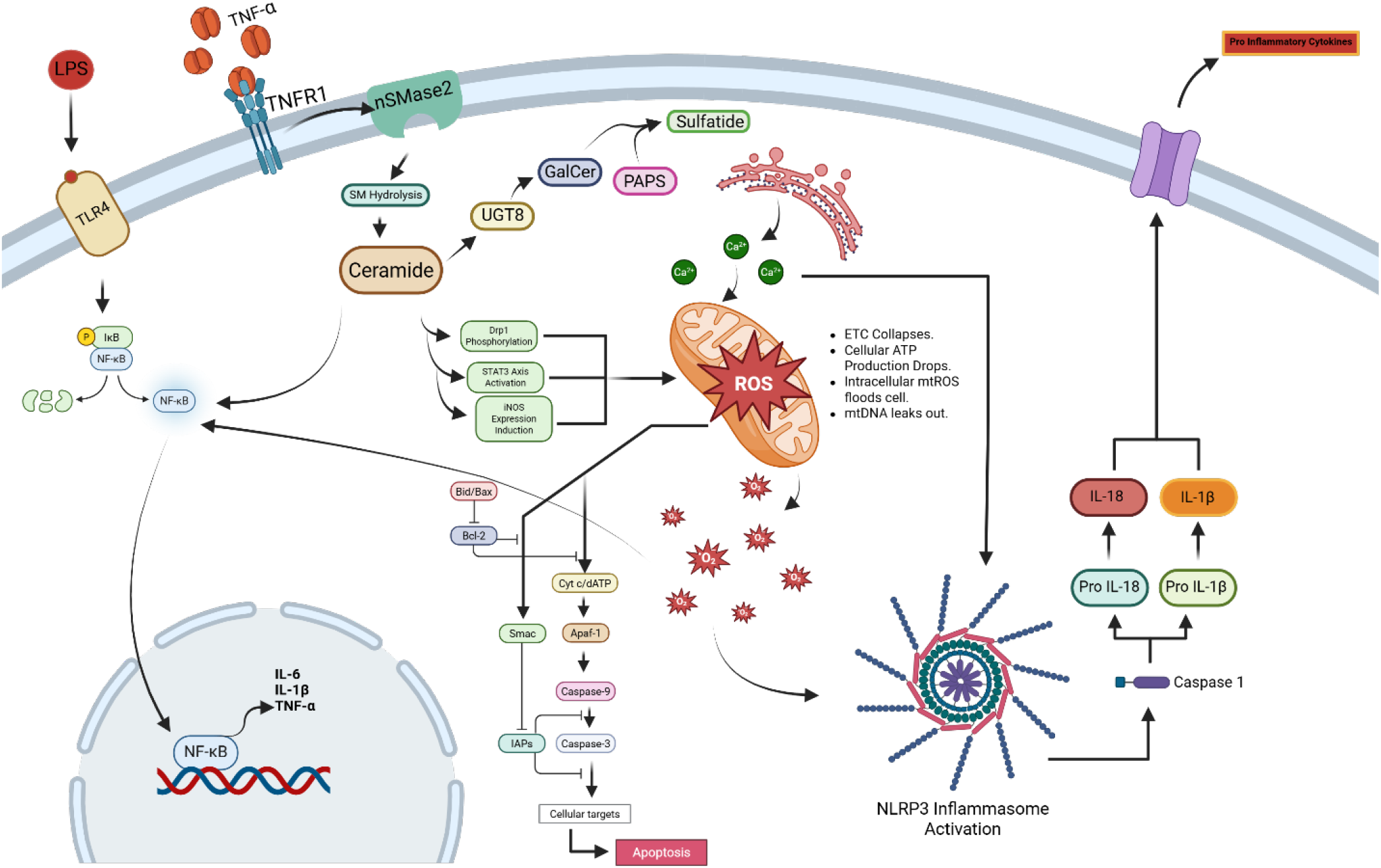
Schematic of the proposed core mechanism linking LPS-induced neuroinflammation to age-dependent sphingolipid remodeling and mitochondrial oxidative stress. LPS activates TLR4 and TNF-α/TNFR1 signaling, leading to nSMase2 activation and hydrolysis of sphingomyelin to generate ceramide. Ceramide is subsequently converted to sulfatide via UGT8, contributing to the sulfatide accumulation detected by MALDI-MSI. In parallel, ceramide promotes mitochondrial ROS production through Drp1 phosphorylation, STAT3 activation, and iNOS induction. Elevated mtROS triggers NLRP3 inflammasome activation, thereby amplifying the inflammatory response. In aged animals, heightened nSMase2 activity is proposed to accelerate this pathway, resulting in more rapid sphingomyelin depletion and greater sulfatide accumulation compared with young animals. This model integrates experimental MALDI-MSI lipidomics findings with established inflammatory and mitochondrial signaling cascades.

### 3.7 Direct comparison of sulfatide distribution between young and old rats

Direct side-by-side comparison of C24 sulfatides (Figure 4) confirmed that old rats exhibited markedly higher signal intensity and broader, less spatially restricted distribution than young rats at both 24 h and 72 h post-LPS. These age-related differences were already detectable at 24 h but became substantially more pronounced at 72 h, particularly for C24(OH)-sulfatide. While young rats maintained relatively structured and regionally confined sulfatide signals even at the later time point, old rats displayed highly diffuse and scattered distribution patterns that extended across large areas of the brain sections. This morphological contrast demonstrates that the age-dependent exacerbation of sulfatide remodeling is not only quantitative but also qualitative, involving a clear loss of spatial organization in aged animals.

These findings indicate that aging alters both the magnitude and the temporal dynamics of sulfatide remodeling during neuroinflammation. The markedly greater differences observed at 72 h compared with 24 h suggest that the effects of aging on this process are cumulative, likely reflecting sustained nSMase2 activity and progressive ceramide production in old rats over time. The particularly strong divergence in C24(OH)-sulfatide distribution at 72 h further supports the idea that longer-chain, hydroxylated sulfatides are preferentially affected in the aged brain under prolonged inflammatory conditions. This species-specific pattern may reflect differences in enzyme substrate preference or greater metabolic stability of hydroxylated species during sustained inflammation.

By enabling direct visual and spatial comparison, Figure 4 highlights that the age-dependent increase in sulfatides is not limited to overall abundance but also involves a fundamental loss of spatial containment. In young rats, sulfatide remodeling remained relatively localized and anatomically organized, whereas in old rats it became diffusely distributed across brain regions. These spatial patterns are consistent with the quantitative elevations detected by peak area analysis (Table 2) and reinforce the conclusion that aging impairs the brain’s ability to regionally restrict and resolve inflammation-driven lipid changes. The progressive nature of this loss of spatial organization from 24 h to 72 h suggests that the aged brain not only initiates a stronger sulfatide response but also fails to effectively contain or reverse these alterations over time, resulting in more extensive and persistent sphingolipid dysregulation, particularly within white matter.

### 3.8 Time-dependent changes in sphingomyelin distribution in young and old rats

MALDI-MSI analysis revealed moderate time-dependent changes in sphingomyelin distribution in young rats following LPS challenge (Supplementary Figure S1, Panels A– F). While SM(d36:1) remained relatively stable in both intensity and spatial distribution between 24 h and 72 h, SM(d18:1/18:0) showed a noticeable increase in signal intensity at the later time point. The distribution of these sphingomyelin species remained relatively structured and regionally defined throughout, with high-intensity areas largely respecting anatomical boundaries. This pattern suggests that young animals undergo limited and spatially organized sphingomyelin remodeling during the resolution phase of inflammation, consistent with effective regulatory mechanisms that prevent excessive or prolonged sphingomyelin hydrolysis.

In old rats, high-quality sphingomyelin images were limited; however, the available sections showed markedly reduced signal intensity and a more fragmented, diffuse distribution compared with young rats, particularly at 72 h post-LPS (Supplementary Figure S1, Panels G and H). This morphological transition from organized to fragmented and less spatially coherent patterns indicates that sphingomyelin depletion in aged animals is accompanied by progressive disruption of normal membrane microdomain organization. The fragmented appearance at the later time point is consistent with sustained rather than transient nSMase2 activity in the aged brain, leading to ongoing sphingomyelin hydrolysis and continued generation of bioactive ceramide.

Although constrained by limited high-quality imaging data in old rats, these spatial patterns align with the quantitative reductions in sphingomyelin levels and, when considered alongside the concurrent accumulation of long-chain sulfatides (Figures 1–4 and Table 2), support a model of age-dependent sphingolipid metabolic rerouting. In young rats, sphingomyelin hydrolysis appears to be transient and spatially contained, allowing for stabilization of lipid distribution by 72 h. In contrast, old rats exhibit cumulative sphingomyelin depletion and spatial fragmentation that worsen between 24 h and 72 h, suggesting that aging impairs the brain’s ability to resolve or spatially restrict inflammation-driven sphingomyelin hydrolysis. This sustained metabolic flux likely contributes to prolonged ceramide production and its subsequent diversion into sulfatide synthesis, thereby exacerbating membrane disruption and white matter vulnerability in the aged brain during neuroinflammation.

### 3.9 Integration of spatial and quantitative MALDI-MSI data

The integration of high-resolution spatial imaging and quantitative peak-area data from MALDI-MSI provided a multidimensional understanding of sphingolipid remodeling that could not be achieved through either approach in isolation. Quantitative measurements (Table 2) demonstrated significant age-related increases in long-chain sulfatides (particularly C24:1-sulfatide and C24(OH)-sulfatide) alongside concurrent reductions in sphingomyelins (especially SM(d36:1)) following LPS challenge. However, spatial imaging revealed that these alterations were not uniformly distributed across the brain. Instead, they exhibited distinct regional, morphological, and temporal patterns that varied markedly between young and old animals.

In young rats, sulfatide signals showed moderate and progressive increases in intensity from 24 h to 72 h while maintaining relatively structured and regionally confined distribution (Figures 2A and 2B). In contrast, old rats exhibited substantially higher signal intensity and a highly diffuse, less anatomically restricted pattern that became especially pronounced at 72 h (Figures 3A, 3B, and 4). This morphological transition from organized to scattered and “cloud-like” distribution was most evident in white matter regions. Concurrently, sphingomyelin signals in old rats appeared fragmented and reduced in intensity at 72 h (Supplementary Figure S1), in stark contrast to the relatively structured distribution maintained in young rats. These spatially resolved patterns align closely with the quantitative data and confirm that the observed lipid changes represent genuine biological remodeling rather than technical artifacts.

Collectively, these integrated findings demonstrate that aging modifies sphingolipid metabolism in a spatially heterogeneous and time-dependent manner during neuroinflammation. The pronounced accumulation of sulfatides in white matter, coupled with the fragmentation and depletion of sphingomyelins in old rats, indicates that myelinated regions undergo more extensive lipid remodeling under inflammatory conditions in the aged brain. This regional vulnerability is consistent with a model in which sustained nSMase2 activity drives enhanced sphingomyelin hydrolysis, leading to elevated ceramide levels that are redirected toward sulfatide synthesis while simultaneously promoting mitochondrial oxidative stress. The progressive nature of these spatially heterogeneous changes from 24 h to 72 h further suggests that aging impairs the brain’s ability to spatially contain and resolve inflammation-driven lipid alterations, resulting in more widespread and persistent disruption of membrane organization and myelin integrity in old animals compared with young ones.

### 3.10 Conceptual kinetic modeling of age-dependent sphingolipid remodeling

To investigate the mechanistic basis of the observed age-dependent sphingolipid changes, a simplified Michaelis-Menten kinetic model was developed to describe the conversion of sphingomyelin to ceramide via nSMase2 and the subsequent formation of sulfatide. The model is defined by the following system of ordinary differential equations:

d[SM]dt =-VmaxSMKm+[SM] d[Cer]dt =VmaxSMKm+[SM]-k2⋅[Cer] d[Sulf]dt =k2⋅[Cer]

where Vmaxrepresents the apparent maximum rate of sphingomyelin hydrolysis, Kmis the Michaelis constant, and k2is the first-order rate constant for ceramide conversion to sulfatide.

The model was used to simulate lipid dynamics over 72 hours under conditions representing young and old rats following LPS challenge. Initial conditions were normalized to baseline MALDI-MSI signals. Parameters were manually tuned as follows: Young rats, Vmax=0.018, k2=0.012; Old rats, Vmax=0.085, k2=0.045; shared Km=0.8. Sulfatide outputs were scaled by a factor of 2.2 to facilitate comparison with relative MALDI-MSI signal intensities.

As shown in Figure S2A, the model predicted substantially faster and more extensive depletion of sphingomyelin in old rats compared with young rats. This was achieved by assigning a significantly higher apparent maximum reaction velocity (Vmax) in the old group, reflecting elevated sphingomyelinase-like activity. The simulated trajectories were qualitatively consistent with normalized experimental MALDI-MSI measurements at 24 h and 72 h, supporting the conclusion that aging is associated with accelerated sphingomyelin hydrolysis during neuroinflammatory responses.

Figure S2B shows the predicted accumulation of sulfatide over time. The model demonstrated substantially greater sulfatide accumulation in old rats than in young rats, with the divergence becoming increasingly evident after 24 h. This age-dependent difference indicates that aging promotes enhanced conversion of sphingomyelin-derived ceramide into sulfatide species, resulting in progressive lipid accumulation that becomes particularly pronounced at later stages of inflammation.

The temporal changes in the Old/Young ratio for sphingomyelin and sulfatide are presented in Figure S2C. The model predicted a progressive divergence between the two age groups over time, with accelerated sphingomyelin depletion and enhanced sulfatide accumulation in aged animals. These ratio trajectories demonstrate that the effects of aging on sphingolipid remodeling are cumulative and become more pronounced with prolonged exposure to inflammatory stimuli, consistent with the experimental observation that the most dramatic differences between young and old rats emerged at 72 h (Figure 4).

Figure S2D illustrates the predicted dynamics of ceramide, the central metabolic intermediate in the model. In old rats, the simulation showed a rapid early rise in ceramide levels followed by a gradual decline as ceramide was converted into sulfatide. In young rats, ceramide levels increased more slowly and remained lower throughout the time course. These dynamics are consistent with enhanced sphingolipid turnover in the aged brain, where increased sphingomyelinase activity generates a transient but significant ceramide surge that is subsequently channeled into downstream pathways, including sulfatide synthesis.

Collectively, these simulations support a mechanism in which aging increases sphingomyelinase-driven lipid remodeling. By assigning a higher Vmax to old rats, the model reproduces the experimental trends of faster sphingomyelin depletion, transient ceramide elevation, and greater sulfatide accumulation observed in the MALDI-MSI data. The results indicate that aging amplifies both the rate and extent of sphingolipid metabolic rerouting, primarily through elevated nSMase2-like activity. This leads to depletion of membrane sphingomyelin, elevated ceramide flux, and progressive accumulation of sulfatide species, particularly in white matter regions. The kinetic modeling thus provides a coherent mechanistic framework that aligns with the quantitative (Table 2), spatial (Figures 1–4, S1), and temporal patterns observed experimentally, and supports the hypothesis that sustained ceramide production contributes to mitochondrial oxidative stress and increased vulnerability of the aged brain to inflammatory insults.

## Discussion

Neutral sphingomyelinase 2 (nSMase2) serves as a central mechanistic link between aging, neuroinflammation, and sphingolipid remodeling in the brain [33,34,35]. Upon LPS exposure, TLR4 and TNF-α signaling rapidly activate nSMase2 to hydrolyze sphingomyelin at the plasma membrane, generating ceramide. In aged animals, both basal and stimulus-induced nSMase2 activity are elevated, leading to accelerated sphingomyelin depletion, particularly of shorter-chain species such as SM(d36:1), and increased ceramide availability. This excess ceramide is preferentially channeled into the sulfatide biosynthetic pathway [36]. Quantitative peak-area analysis (Table 2) and high-resolution spatial MALDI-MSI (Figures 1-4) demonstrate that this age-dependent metabolic rerouting is not only quantitatively greater but also spatially disorganized and temporally progressive in old rats compared with young animals.

Quantitative data revealed a clear divergence between lipid classes. Old rats exhibited a ∼2.12-fold increase in C24:1-sulfatide and a ∼1.45-fold increase in C24(OH)-sulfatide, accompanied by an approximately 10-fold reduction in SM(d36:1). These opposing changes were substantially amplified at 72 h compared with 24 h, indicating that the effects of aging on sphingolipid metabolism are cumulative. Spatial imaging further revealed that sulfatide accumulation in old rats became highly diffuse and less anatomically restricted by 72 h (Figures 3B and 4), particularly in white matter regions such as the corpus callosum, while sphingomyelin signals appeared fragmented and reduced in intensity (Supplementary Figure S1). In contrast, young rats displayed moderate, progressive, and spatially contained increases in sulfatide intensity with relatively preserved sphingomyelin organization (Figures 2A and 2B). This morphological contrast demonstrates that aging impairs the brain’s capacity to spatially contain and resolve inflammation-driven lipid remodeling.

While nSMase2-mediated sphingomyelin hydrolysis represents the dominant route to ceramide, additional contributions can arise from de novo synthesis, sphingosine salvage/recycling, and glucosylceramide or ceramide-1-phosphate intermediates (Supplementary Figures S3-S5) [37,38,39,40,41,42]. In the aged brain under inflammatory challenge, metabolic flux is preferentially redirected toward sulfatide accumulation. This coordinated rerouting is supported by the inverse relationship between sphingomyelin depletion and sulfatide accumulation, as well as the preferential increase in longer-chain and hydroxylated sulfatide species. These changes align with the substrate preferences of oligodendrocyte-enriched CerS2 and the inflammation-induced upregulation of FA2H and CST, as detailed in the biosynthetic pathway schematics (Supplementary Figures S6-S11).

Kinetic modeling provided mechanistic support for these experimental observations. A simplified Michaelis-Menten model, parameterized with a significantly higher apparent maximum reaction velocity in old rats, qualitatively reproduced the faster sphingomyelin depletion, transient ceramide surge, and greater sulfatide accumulation observed in the MALDI-MSI data (Supplementary Figure S2). Parameter sensitivity analysis confirmed that model outputs were most sensitive to increases in the nSMase2-related rate constant, underscoring this enzyme as the primary driver of age-dependent remodeling.

Ceramide generated by nSMase2 occupies a central position that drives a self-reinforcing cycle. While a portion of ceramide is converted into sulfatide via the CerS2-FA2H-UGT8-CST cascade, excess ceramide promotes Drp1 recruitment to mitochondria, triggering excessive fission, elevating mitochondrial ROS production, and impairing bioenergetics [43,44]. This oxidative stress and pro-inflammatory signaling, in turn, prolong nSMase2 activation and further upregulate downstream enzymes in oligodendrocytes, generating additional ceramide and perpetuating the loop [45,41]. In young rats, this cycle remains transient and spatially contained, allowing sphingomyelin distribution to stabilize by 72 h. In old rats, however, the cycle becomes self-perpetuating due to sustained nSMase2 activity, age-related impairment of arylsulfatase A-mediated sulfatide degradation, and reduced resolution capacity associated with microglial senescence. Consequently, old animals exhibit cumulative sulfatide accumulation, particularly of myelin-enriched C24 and hydroxy-C24 species, and progressive sphingomyelin depletion and fragmentation, with increasingly diffuse spatial patterns, especially in white matter.

These lipid alterations carry significant biological consequences. Excessive long-chain and hydroxylated sulfatide accumulation can alter myelin lipid packing, increase membrane rigidity, and disrupt lipid-raft integrity normally supported by CerS2-derived very-long-chain sphingolipids. Concurrent sphingomyelin depletion further compromises raft organization and membrane signaling. Most critically, the transient but significant ceramide surge directly targets mitochondria, promoting Drp1-mediated fission and elevated mtROS that disproportionately affect oligodendrocytes and axons, cells with high metabolic demand [45,48]. Inflammation-driven upregulation of FA2H and CST, combined with sustained ceramide supply, leads to overproduction and mislocalized accumulation of myelin-enriched sulfatides, further compromising myelin stability and remyelination capacity. These alterations are therefore positioned to contribute to the white-matter dysfunction commonly observed in aging and neurodegenerative diseases [44].

The therapeutic implications of these findings are considerable. Because nSMase2 acts as a key driver of this self-reinforcing ceramide-mitochondrial cycle, targeted inhibition represents a promising strategy to interrupt upstream ceramide generation, limit substrate availability for the upregulated CerS2-FA2H-CST axis, and reduce downstream mitochondrial oxidative stress. Pharmacological nSMase2 inhibitors have shown efficacy in preclinical neuroinflammation models and could be evaluated in age-related contexts. Combination approaches, pairing nSMase2 inhibition with mitochondrial protectants (e.g., Drp1 modulators) or senolytics to clear senescent microglia, may offer synergistic benefits by addressing both lipid metabolic and cellular senescence components of inflammaging. Additionally, engineered extracellular vesicles delivered intranasally provide a feasible route for CNS-targeted delivery of miRNAs or small-molecule inhibitors. Beyond therapy, the distinct molecular species and progressive white-matter accumulation patterns identified here may serve as candidate biomarkers for monitoring neuroinflammatory burden and white-matter vulnerability in aging and neurodegenerative disease.

Collectively, this study demonstrates that aging fundamentally reprograms sphingolipid metabolism during neuroinflammation through sustained nSMase2 activity, resulting in progressive, spatially diffuse accumulation of long-chain and hydroxylated sulfatides and sphingomyelin depletion that converge on mitochondrial oxidative stress and white-matter vulnerability. The integration of high-resolution MALDI-MSI, quantitative analysis, kinetic modeling, and biosynthetic pathway mapping provides a powerful framework for identifying the nSMase2-ceramide-CerS2/FA2H/CST axis as a modifiable mechanism linking neuroinflammaging, oligodendrocyte dysfunction, and brain aging.

## Conclusion

This work reveals that aging amplifies and spatially reorganizes sphingolipid remodeling during LPS-induced neuroinflammation. Old rats displayed markedly accelerated sphingomyelin depletion and progressive, diffuse accumulation of long-chain and hydroxylated sulfatides, particularly C24 and C22 species in white matter, in contrast to the more controlled and spatially contained response in young animals. These changes were driven by sustained nSMase2 activity that elevates ceramide, which is then routed through upregulated oligodendrocyte enzymes (CerS2, FA2H, UGT8, CST) to produce specific molecular species. The integration of spatial imaging, quantitative data, kinetic modeling, and pathway analysis provides strong evidence that aging impairs the brain’s ability to resolve inflammation-driven lipid alterations, resulting in cumulative metabolic and mitochondrial dysfunction with direct consequences for myelin integrity. These findings position the nSMase2-ceramide-CerS2/FA2H/CST pathway as a promising therapeutic target for mitigating age-related white-matter vulnerability and neuroinflammatory damage.

## Limitations

Several limitations should be acknowledged. First, the kinetic model was conceptual; parameters were manually tuned rather than formally optimized against individual replicates. Future studies with direct measurements of regional nSMase2, CerS2, FA2H, and CST activity will be needed to refine parameter estimation. Second, high-quality sphingomyelin imaging data from old rats were limited. Third, direct measurements of mitochondrial ROS, Drp1 activation, and bioenergetic function were not performed. Fourth, a single acute ICV LPS stimulus may not fully capture chronic or sterile neuroinflammation. Finally, functional outcomes such as myelin ultrastructure, axonal conduction, and cognitive performance were not assessed. Future investigations incorporating these measurements, along with larger sample sizes and additional aging/inflammation models, will be essential to strengthen and extend the proposed framework.

## ACKNOWLEDGMENT

We sincerely acknowledge financial support from the NIH (R01HL163159, Z.S.; R15 EB035866, L.B.), the American Heart Association (AHA grant #1807047, L.B.), and GLRC-ICC (R01805, L.B.). All authors, especially the graduate students, gratefully acknowledge the Department, Dr. Will Cantrell, and the Graduate School for their support of student training and research progress. We also extend our deep gratitude to Dr. Rick Koubek for his encouragement and unwavering support throughout this project.

## Abbreviations

AD: Alzheimer’s disease
C1P: ceramide-1-phosphate
CerS: ceramide synthase
CNS: central nervous system
DAMPs: damage-associated molecular patterns
GalCer: galactosylceramide
ICV: intracerebro-ventricular
LPS: lipopolysaccharide
MALDI-MSI: matrix-assisted laser desorption/ionization mass spectrometry imaging
nSMase 2: neutral sphingomyelinase 2
PD: Parkinson’s disease
S1P: sphingosine-1-phosphate
SASP: senescence-associated secretory phenotype
TLR4: toll-like receptor 4
Vmax: maximum reaction velocity.

