## Supplemental Material for "Age-Dependent Spatiotemporal Remodeling of Brain Sphingolipids During LPS-Induced Neuroinflammation: MALDI-MSI Reveals Accelerated Sphingomyelin Depletion and Sulfatide Accumulation Linked to Mitochondrial Oxidative Stress"

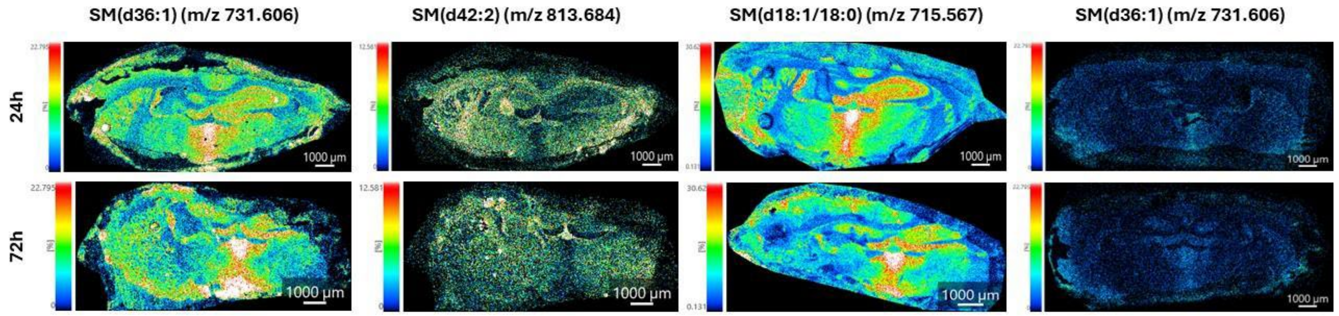


Figure S1. Spatial distribution of sphingomyelin species in young and old rat brains following LPS-induced neuroinflammation. Representative MALDI-MSI images showing the spatial distribution of sphingomyelin species in coronal brain sections from young and old rats at 24 h and 72 h after intracerebroventricular LPS administration. Ion images are shown for SM(d36:1) (m/z 731.606), SM(d42:2) (m/z 813.684), and SM(d18:1/18:0) (m/z 715.567). Color scale bars indicate relative ion intensity (blue = low; red/white = high). Scale bar = 1000 µm. In young rats, sphingomyelin signals remained relatively structured and regionally defined at both 24 h and 72 h, with high-intensity areas largely respecting anatomical boundaries. In contrast, old rats exhibited markedly reduced overall signal intensity and a more fragmented, diffuse distribution, particularly evident at 72 h post-LPS. This morphological transition from organized to fragmented patterns indicates that sphingomyelin depletion in aged animals is accompanied by disruption of normal membrane organization during prolonged neuroinflammation. These spatial alterations are consistent with the quantitative reductions in sphingomyelin abundance observed in aged animals and support enhanced sphingomyelin hydrolysis in the aged brain under inflammatory conditions.

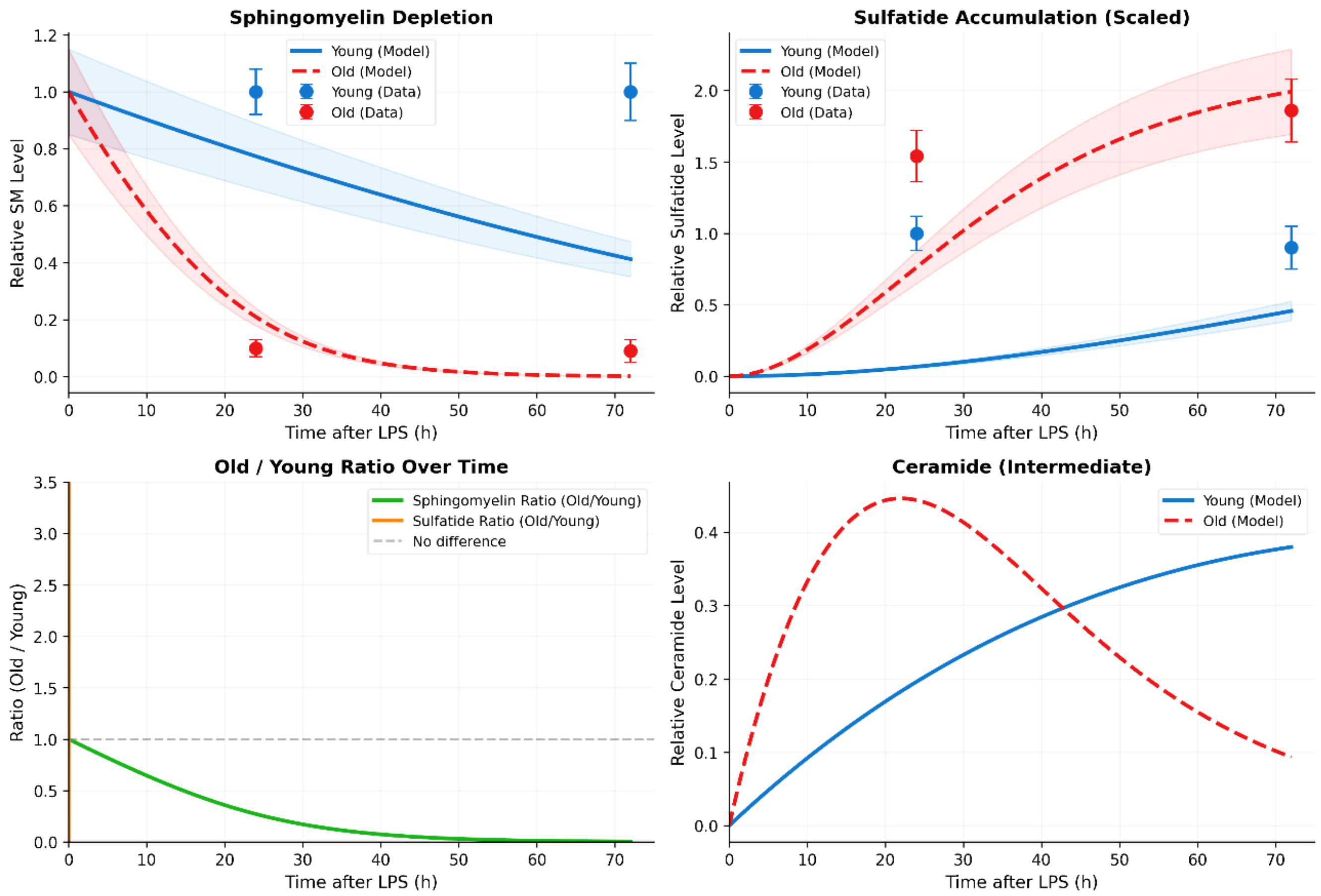


Figure S2. Conceptual Michaelis-Menten model of age-dependent sphingolipid remodeling following LPS-induced neuroinflammation. A simplified kinetic model was constructed to describe the conversion of sphingomyelin to ceramide and subsequent formation of sulfatide using Michaelis-Menten kinetics. Model parameters were manually tuned to approximate experimental trends observed in MALDI-MSI datasets. Sulfatide outputs were scaled by a factor of 2.2 to account for differences between model-predicted concentrations and relative MALDI-MSI signal intensities. Shaded regions represent ±15% model uncertainty. Symbols with error bars represent normalized experimental MALDI-MSI measurements at 24 h and 72 h post-LPS. (A) Predicted sphingomyelin (SM) depletion in young and old rats. The model predicts substantially faster and more extensive depletion of sphingomyelin in old rats compared with young rats, consistent with elevated sphingomyelinase-like activity in aged animals. (B) Predicted sulfatide accumulation over time. The model predicts markedly greater sulfatide accumulation in old rats than in young rats, consistent with experimental observations of elevated C24 sulfatide levels in aged animals at 72 h post-LPS. (C) Temporal changes in the Old/Young ratio for sphingomyelin and sulfatide. The model predicts progressive divergence between age groups over time, with accelerated sphingomyelin depletion and enhanced sulfatide accumulation in old rats. The dashed horizontal line indicates no age-related difference (ratio = 1). (D) Predicted ceramide dynamics. The model illustrates ceramide as a transient intermediate, showing a rapid early surge in old rats followed by a gradual decline, consistent with enhanced sphingolipid turnover under conditions of elevated sphingomyelinase activity. Model parameters: Young rats, Vmax = 0.018, $k_2$ = 0.012; Old rats, Vmax = 0.085, $k_2$ = 0.045; Km = 0.8.

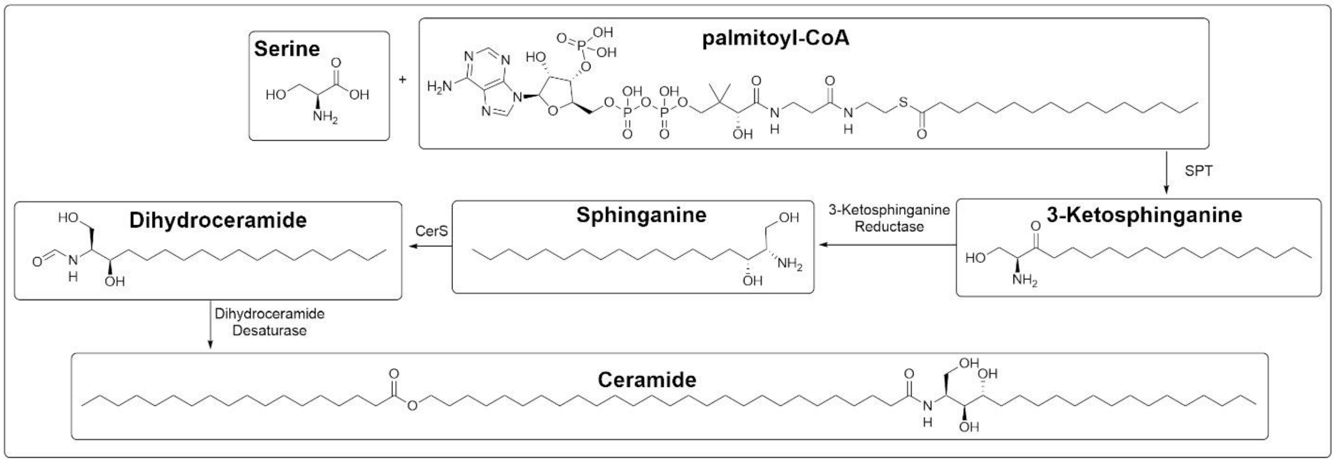


Figure S3. De novo ceramide biosynthesis pathway. Schematic representation of the de novo biosynthetic route for ceramide production. Serine and palmitoyl-CoA are condensed by serine palmitoyltransferase to form 3-ketosphinganine. This intermediate is reduced by 3-ketosphinganine reductase to sphinganine, which is then N-acylated by ceramide synthases (CerS) to generate dihydroceramide. Finally, dihydroceramide desaturase introduces a double bond to produce ceramide. This pathway provides an alternative source of ceramide in addition to nSMase2-mediated hydrolysis of sphingomyelin and may contribute to the elevated ceramide flux observed in aged animals during neuroinflammation.

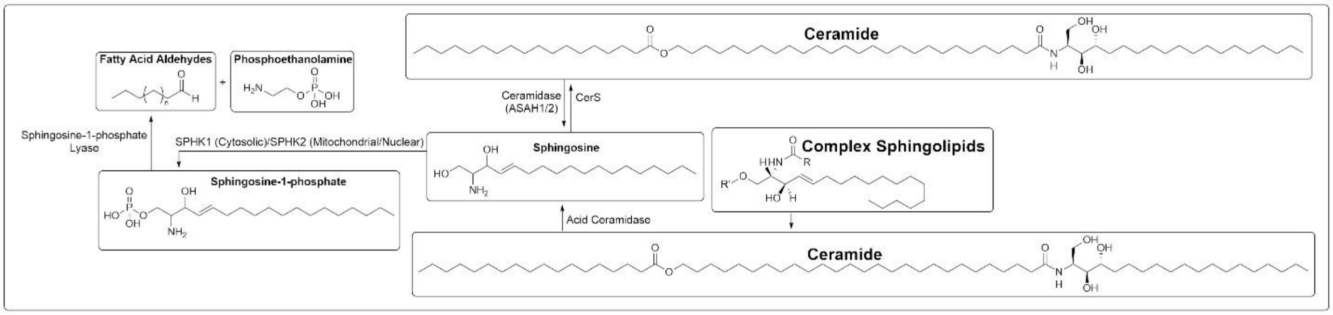


Figure S4. Ceramide salvage pathway and sphingosine-1-phosphate (S1P) signaling. Schematic representation of the ceramide salvage (recycling) pathway and S1P metabolism. Ceramide can be hydrolyzed by acid ceramidase (ASAH1/2) to generate sphingosine. Sphingosine is then phosphorylated by sphingosine kinases (SPHK1 in the cytosol/lysosomes or SPHK2 in mitochondria/nucleus) to form S1P. S1P can be dephosphorylated back to sphingosine or cleaved by S1P lyase into fatty acid aldehydes and phosphoethanolamine. Ceramide can also be regenerated from sphingosine through the action of CerS. This salvage pathway allows recycling of sphingosine back into ceramide or complex sphingolipids and represents an additional route contributing to the cellular ceramide pool during neuroinflammation.

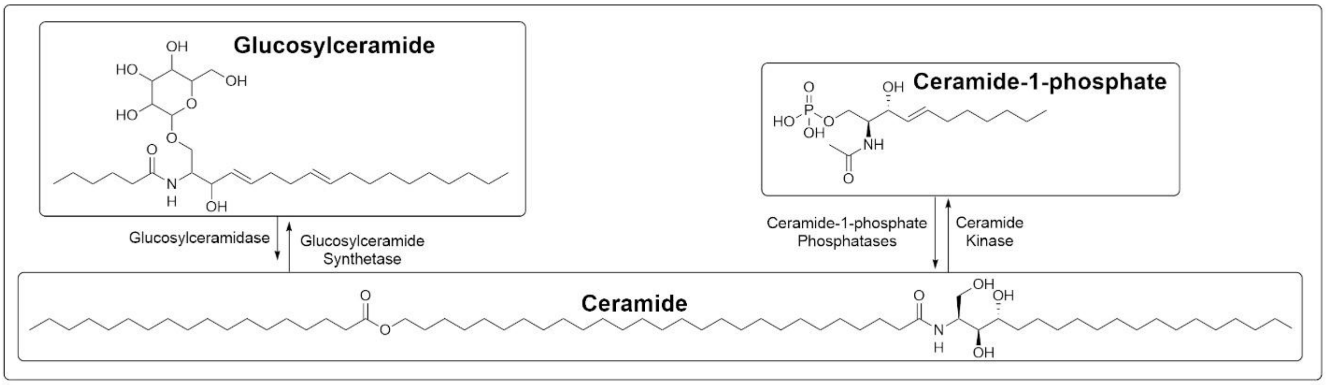

Figure S5. Glucosylceramide and ceramide-1-phosphate (C1P) metabolic pathways. Schematic representation of two alternative routes of ceramide metabolism. On the left, ceramide can be converted to glucosylceramide by glucosylceramide synthase and subsequently hydrolyzed back to ceramide by glucosylceramidase. On the right, ceramide is phosphorylated by ceramide kinase to form C1P, which can be dephosphorylated back to ceramide by C1P phosphatases. These pathways represent additional routes of ceramide generation and consumption that may contribute to sphingolipid remodeling and bioactive lipid signaling during neuroinflammation.

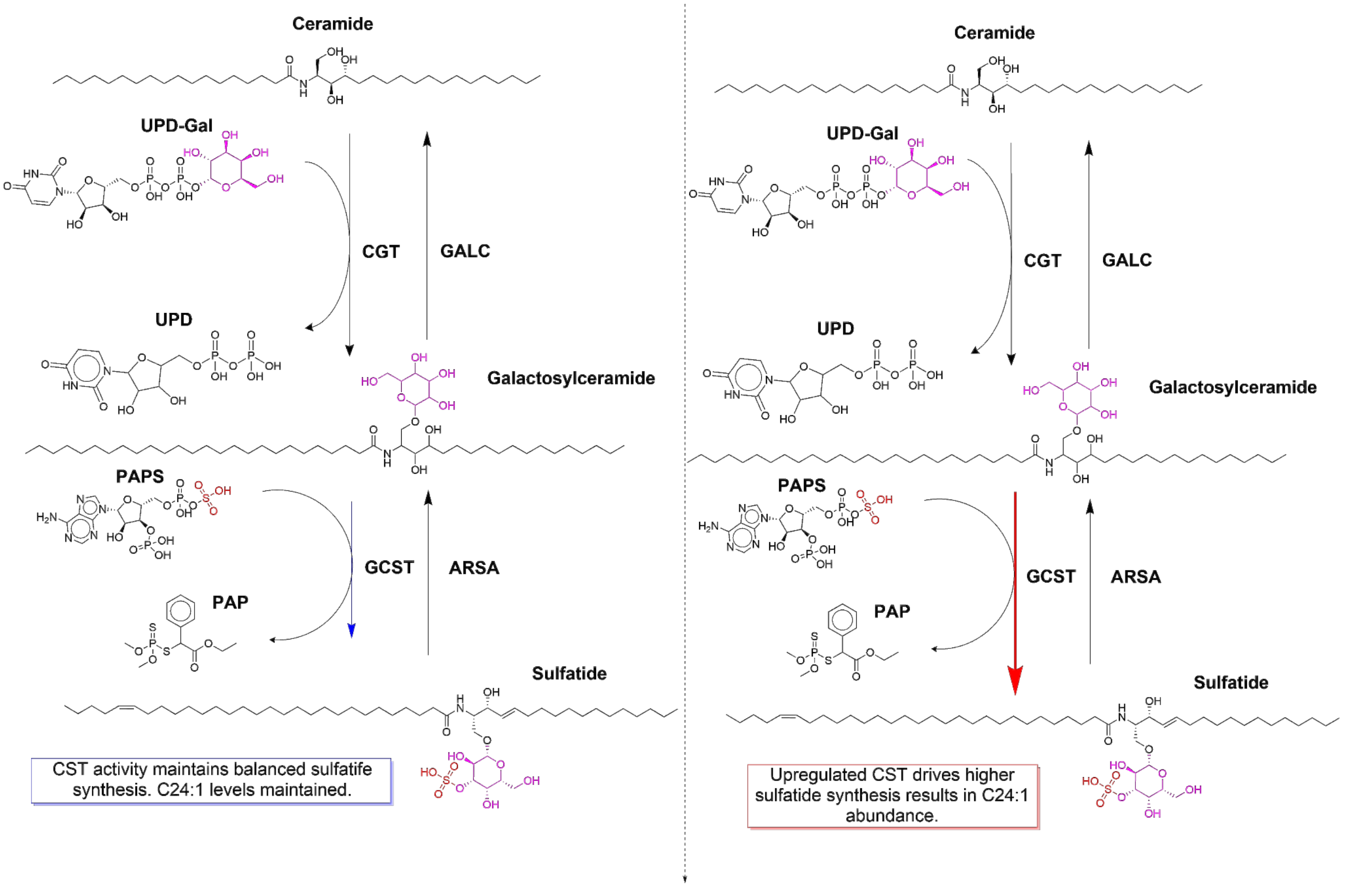


Figure S6. Biosynthesis of C24:1-sulfatide and age-dependent remodeling in LPS-induced neuroinflammation. Comparison of young (left) vs old (right) rats showing ER (CerS2/UGT8) and Golgi (CST/GAL3ST1) steps leading to C24:1-sulfatide (m/z 888.624). Old rats exhibit upregulated CST activity, diffuse white-matter accumulation, and myelin disruption at 24–72 h post-ICV LPS (MALDI-MSI).

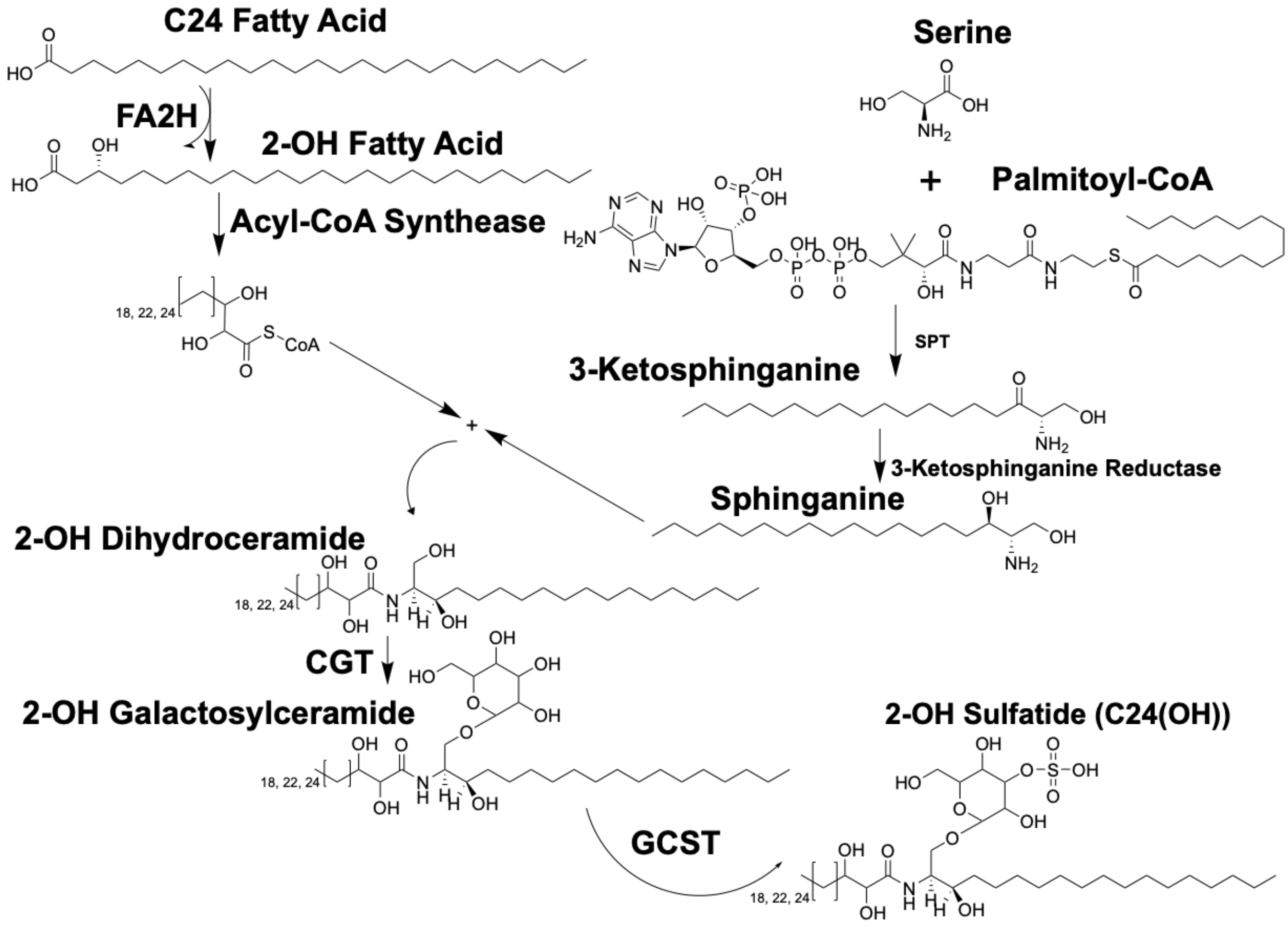


Figure S7. Biosynthesis of C24(OH)-sulfatide and age-dependent remodeling after LPS-induced neuroinflammation. ER (FA2H hydroxylation + CerS2) and Golgi (CST) pathway for hydroxy-C24 sulfatide (m/z 906.635). Young rats maintain structured compact myelin; old rats show upregulated FA2H/CST, highly diffuse “cloud-like” distribution in white matter, and severe myelin disruption at 24–72 h (MALDI-MSI).

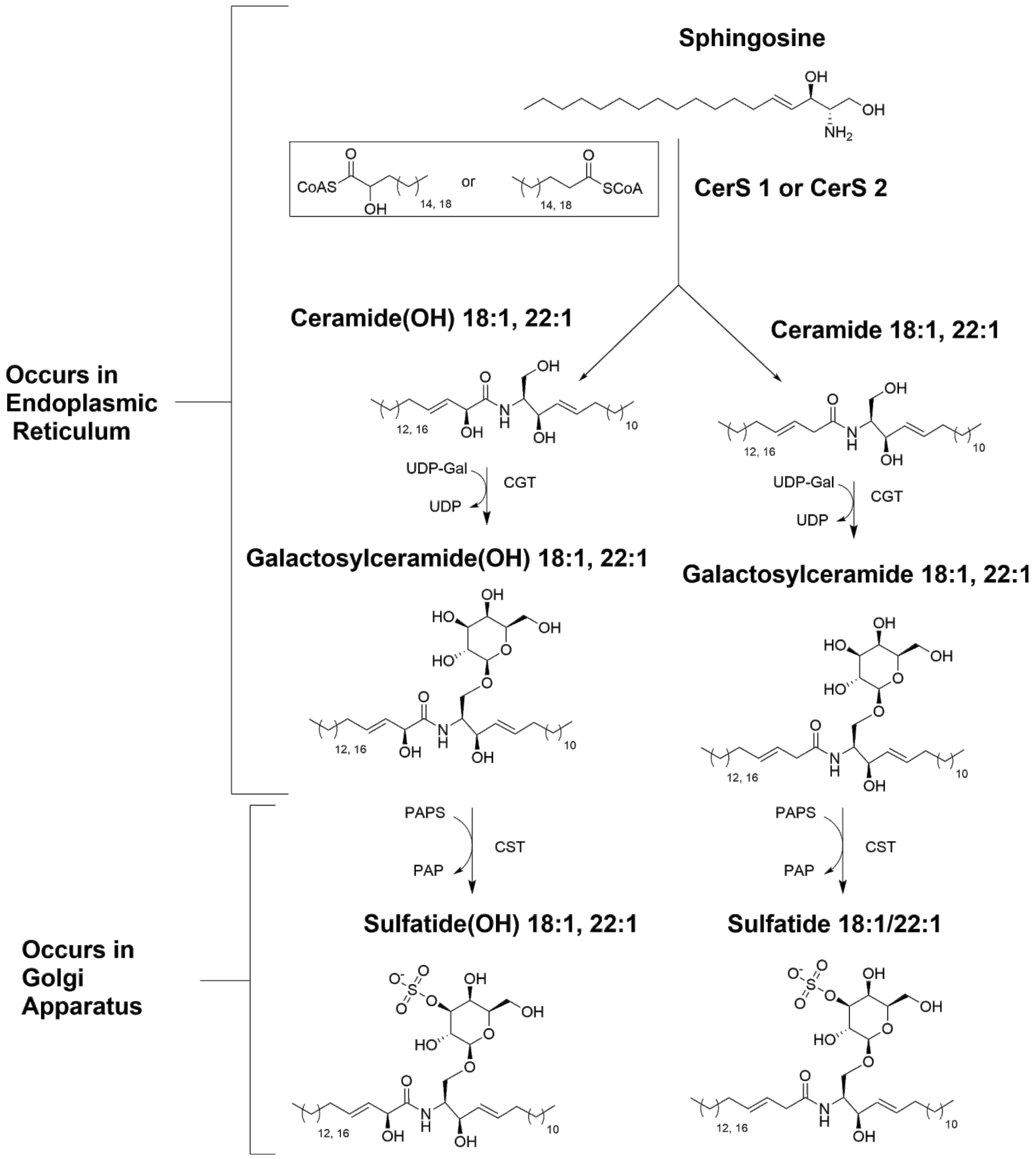


Figure S8. Biosynthesis of C18- and C22-sulfatide species and age-dependent remodeling after LPS neuroinflammation. CerS1/CerS2-driven pathways for C18 (m/z 806.546/822.541) and C22 (m/z 862.608/878.603) sulfatides. Young rats show controlled, regionally confined increases; old rats display exaggerated intensity, diffuse scattered distribution, and loss of anatomical restriction at 24–72 h post-LPS (MALDI-MSI).

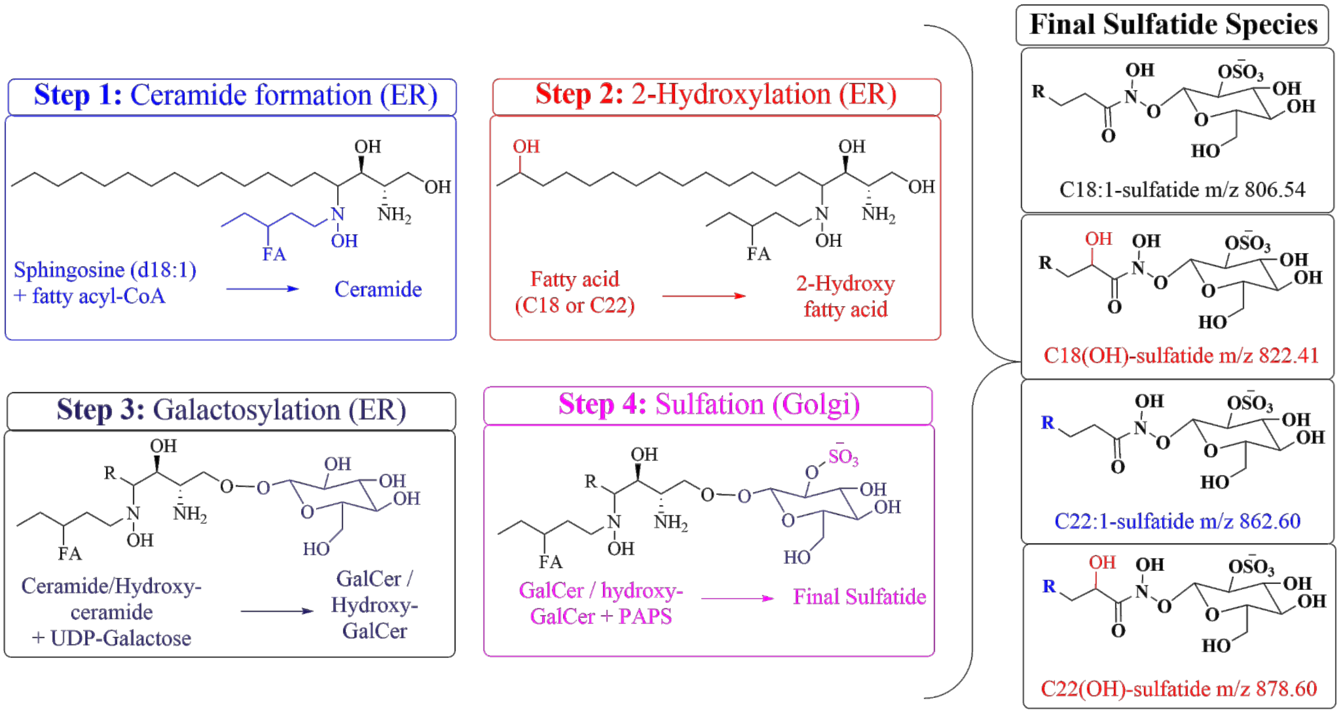


Figure S9. Biosynthesis of C18- and C22-sulfatide species and exaggerated remodeling in old rat brain after LPS neuroinflammation. Stepwise ER-to-Golgi pathway (CerS → FA2H → UGT8 → CST) for C18/C22 and hydroxy species. Old rats exhibit robust progressive upregulation of FA2H/CST, widespread diffuse sulfatide signals (especially white matter), and loss of structured distribution compared with young rats (MALDI-MSI, 24–72 h).

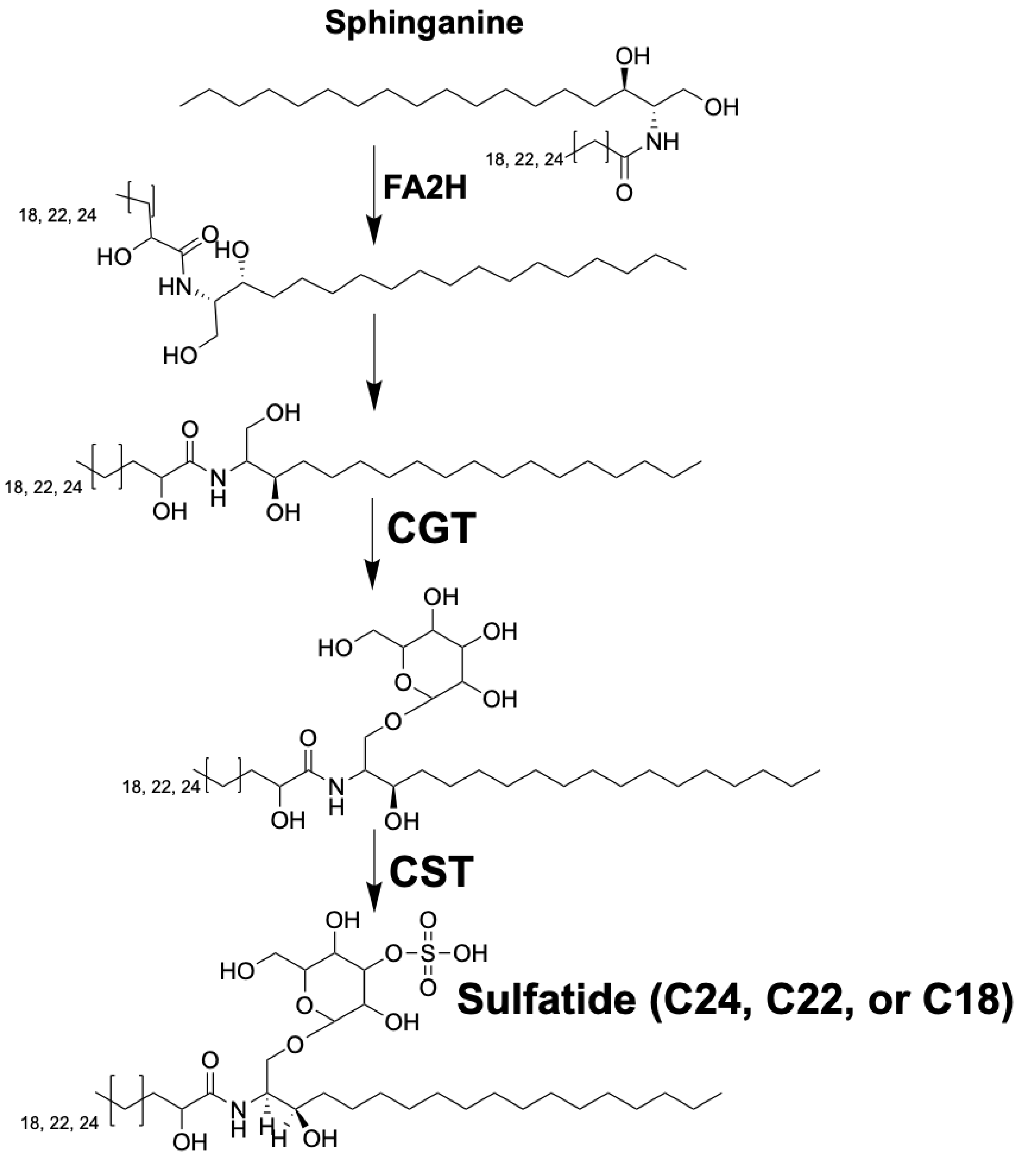


Figure S10. Biosynthesis of myelin-enriched C24 sulfatide species and exaggerated diffuse remodeling in old rat brain after LPS challenge. CerS2/FA2H/UGT8/CST pathway for VLC C24 and hydroxy-C24 sulfatides (m/z 888.624/906.635). Old rats show markedly higher intensity, highly diffuse "cloud-like" white-matter distribution, and progressive spread from 24 h to 72 h; young rats maintain structured, regionally confined patterns (MALDI-MSI).

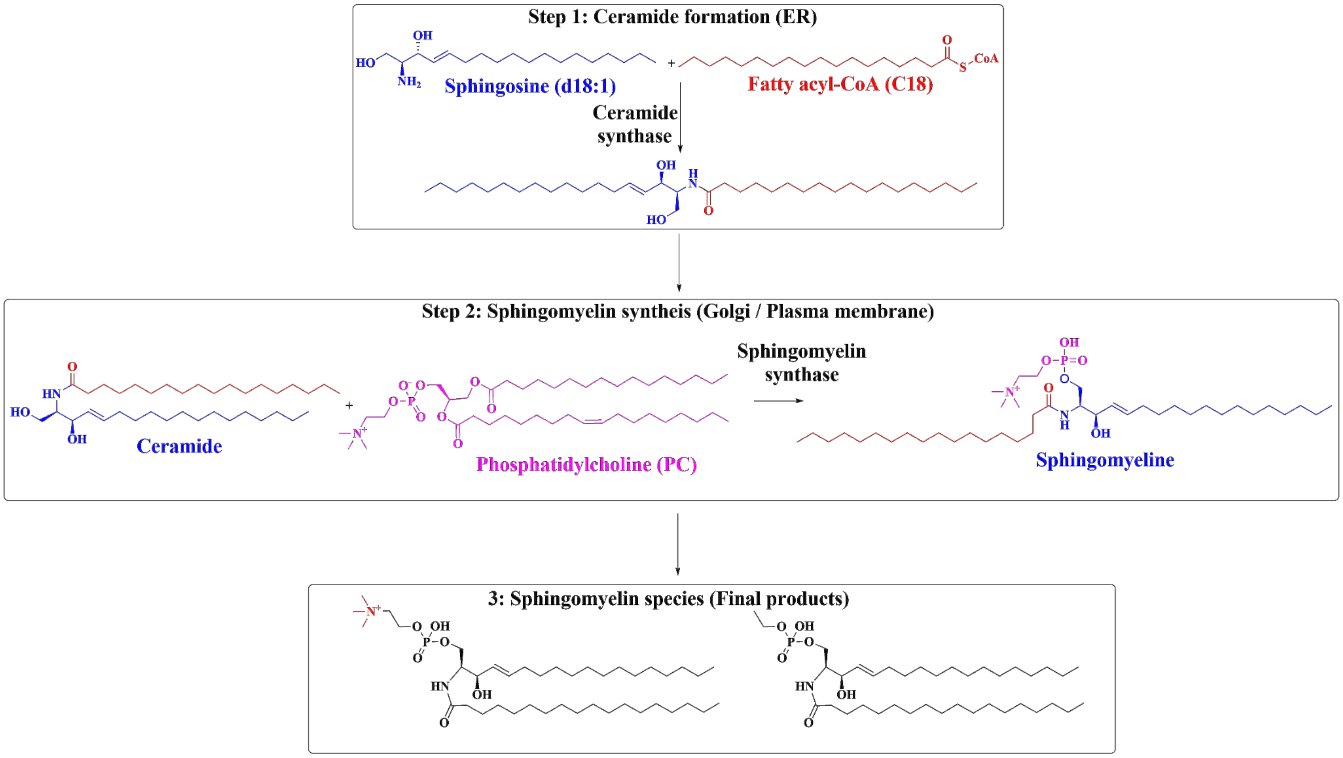


Figure S11. Biosynthesis of sphingomyelin species and age-dependent depletion/fragmentation in rat brain after LPS-induced neuroinflammation. Ceramide → sphingomyelin (SMS1/2) pathway and acid/neutral SMase-mediated hydrolysis. Young rats preserve structured, regionally defined SM distribution; old rats exhibit markedly reduced intensity, highly fragmented/diffuse pattern, and progressive membrane disruption at 24–72 h (MALDI-MSI).
